# Enhanced *Bacillus subtilis* natural competence enables multiplexed genome and spore engineering

**DOI:** 10.64898/2026.05.01.722266

**Authors:** Jessica A. Lee, Nikhil U. Nair

## Abstract

*Bacillus subtilis* is an important chassis for biotechnology, but its use in multiplex genome engineering is limited by low natural transformation efficiency. Here, we compared inducible promoter systems for synthetic activation of the competence regulator ComK and evaluated their effects on the *comG* operon competence reporter and transformation efficiency. Xylose- and mannitol-inducible systems outperformed IPTG-based constructs and shifted 96–99% of cells into a reporter-positive competent state. However, reporter activation alone did not predict transformation potential. Optimization of culture density and induction timing increased transformant yield 45-fold relative to the initial protocol and 2800-fold relative to the conventional Spizizen method. Disruption of native competence regulatory genes did not improve performance and often reduced transformation output, highlighting the importance of endogenous regulatory circuitry. Using the optimized strain and protocol, we achieved co-transformation frequencies of 11–18% and constructed multiplex spore-display libraries containing fluorescent protein fusions integrated at multiple loci. Screening identified strong dual-display combinations and showed that cargo loading depends on anchor protein, integration locus, and genetic background. SscA fusions supported the highest display capacity and promoted synergistic co-display. Together, these results show improvements in natural transformation-based genome engineering in *B. subtilis* and provide insight into the construction of multifunctional engineered spores.

## INTRODUCTION

*Bacillus subtilis* is a Gram-positive, soil-dwelling bacterium which has long served as a model-organism for studying cellular differentiation in bacteria. As *B. subtilis* cells reach stationary phase, complex, interdependent regulatory networks govern differentiation into a variety of cell fates including matrix formation, sporulation, and competency^1^. While *B. subtilis* is currently utilized in industrial production of recombinant proteins, antibiotics, and chemicals, engineering in this bacterium is hindered by comparatively low transformation efficiencies (10^9^ transformants μg^-1^ DNA in *E. coli* vs. 10^6^ transformants μg^-1^ in *B. subtilis*).

*B. subtilis* displays natural competence—the ability to actively uptake exogenous DNA and, given sufficient homology, recombine it into the genome. Natural competence can be a simple, efficient method of genome engineering; in *Vibrio* spp., multiplex genome editing by natural transformation (MuGENT) can produce scarless genome editing close at to 50 % efficiency^2,3^. In *B. subtilis*, genome engineering by natural transformation has been used for the production of genome-scale knock-out libraries and multiplexed genome editing for metabolic engineering applications – albeit at a far lower efficiency of <5 % for distant loci^4,5^ using the Spizizen method^6^. This method activates the natural competence phenotype by growing cells to early stationary phase in a glucose-based minimal media, triggering nutrient limitations and quorum sensing responses. However, even under optimized conditions, only 5−20 % of the population enters the competent state^7^. Activation of promoter of the late competence operon, *comG*, is commonly used to define the competent state. Extensive study of the competence regulatory network has elucidated complex dynamics between hundreds of proteins driving the observed stochasticity in cell fate determination. ComK is the global regulator of competency; several negative regulators keep basal levels of ComK low during exponential growth. However, upon sufficient stress and quorum sensing inputs, ComK levels rise towards an autocatalytic threshold over which only a small portion of the population will cross and become competent. To simplify induction of the competent state, several groups have placed ComK under the control of an inducible promoter, allowing for the induction of competence in rich media^8–10^. Additionally, the *comG* promoter is commonly used as a reporter for competency; but there is little data to show how reporter activity corresponds to transformability of a given cell.

Here, we compare the synthetic induction of ComK under five commonly utilized promoters in *B. subtilis*. Through characterization of these systems, we find the importance of sufficient cell density on transformation frequency. Furthermore, we uncover a complex relationship between reporter activity and transformation frequency—influenced both by growth conditions and interactions between ComK induced from its synthetic and the native loci. The competence reporter shows induction of ComK shifts the entire population into the competent state; however, transformation frequency is still relatively low – only 1 in 10,000. Nevertheless, we selected an inducible competence construct and optimized the protocol to maximize number of transformants—demonstrating a 2800-fold improvement. We then leverage higher transformation and co-transformation frequencies to screen co-transformed multiplexed genomic integrations of spore protein-fluorescent protein fusion constructs. We find several configurations that confer high display of YFP and mCherry on spores and highlight the dependencies on locus, spore protein, and genetic background for display of two proteins.

## RESULTS

### Inducible ComK expression increases the fraction of cells in competence state

Several inducible promoters have been used in *B. subtilis* to induce ComK expression^8–10^. While all inducible systems produce transformants, there is variability in reported transformation efficiencies. To more directly compare relative expression levels between inducible promoters, we built YFP reporter strains for the following five promoters: P_xyl_, P_xyl47_, P_mtl_, P_hyperspank_ (P_hyp_), and P_T7_. P_xyl_, P_mtl_, and P_hyp_ have previously been used to induce competence; however, P_xyl47_ and P_T7_ have not. P_xyl47_ is a modified version of the original *xylA* promoter from the pAX01 plasmid; vestigial portions of the promoter were removed by Castillo-Hair et. al resulting in higher expression^11^. The construct design of P_T7_ follows that of Castillo-Hair et al. which was predicted to have higher expression than 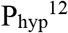. We integrated all constructs at the *lacA* locus. We recorded the response curves for each strain to understand the dynamic range of each promoter (**Figure 1a– e**). Most constructs display Hill-function like behavior—except *P*_*mtl*_-*YFP* which shows maximum expression at intermediate mannitol concentrations. As expected, the *P*_*xyl47*_-*YFP* and *P*_*T7*_-*YFP* constructs display higher fluorescence than the other three constructs (**Figure 1a–e**). P_xyl_, P_mtl_, and P_hyp_ have similar levels of expression. At high concentration of inducer (1 % xylose, 0.5 % mannitol, or 500 μM IPTG), all YFP constructs show unimodal population distributions when analyzed by flow cytometry, indicating uniform induction across the entire population (**Figure 1f**).

**Figure 1.**
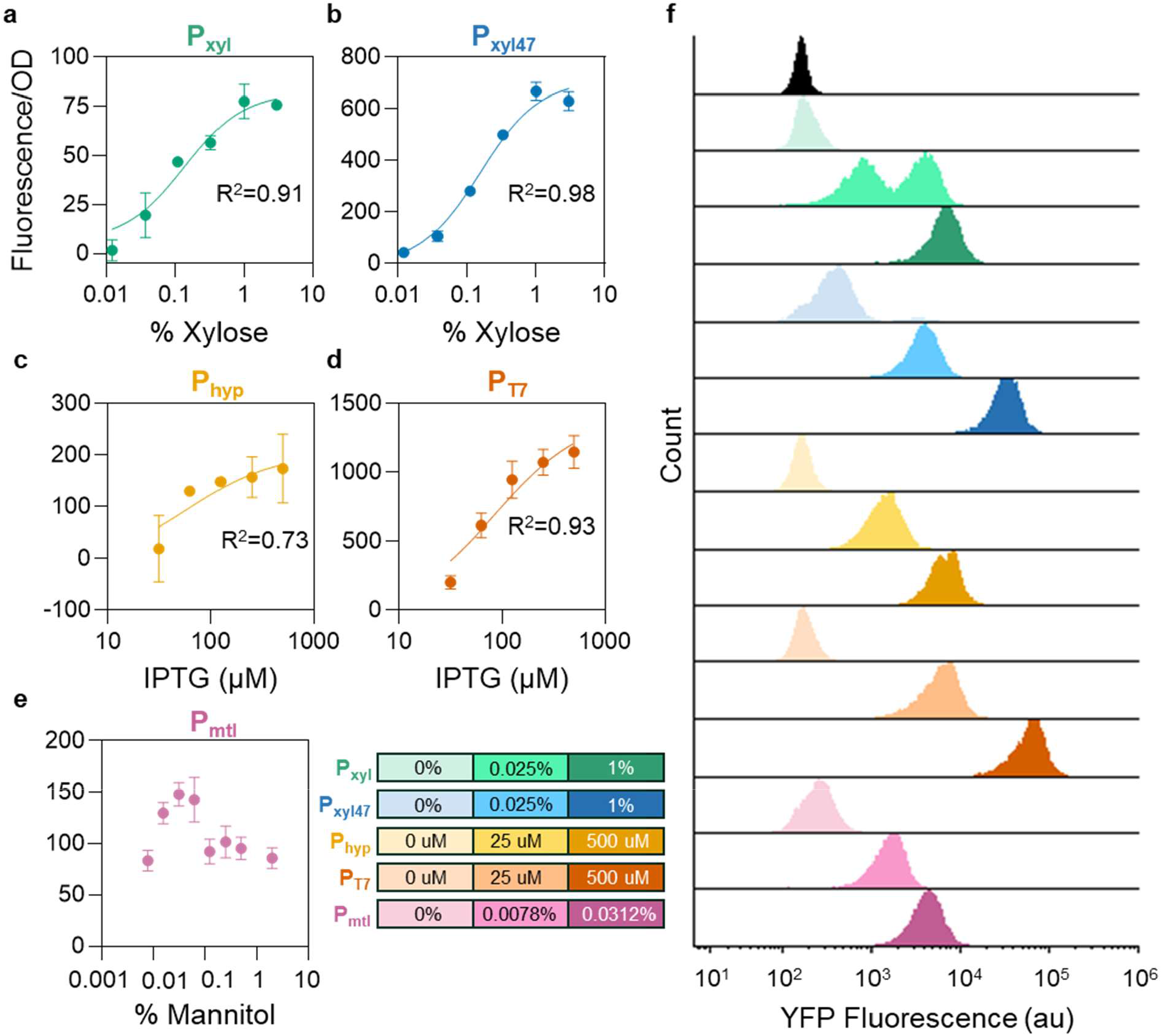
Characterization of common inducible promoters in *B. subtilis*. **a)** Dose response curves of P_*xyl*_-, **b)** P_*xyl47*_-, **c)** P_*hyperspank*_-, **d)** P_*T7*_-, and **e)** P_*mtl*_-YFP. Measurements were taken 2 h post-induction. The data was fitted using Graphpad Prism [Agonist] vs response model. **f)** YFP fluorescence population distribution of each inducible promoter at three concentrations of inducer after 2 h post-induction. Counts are normalized to 100.

To compare existing inducible competence constructs, we followed the published designs for *P*_*xyl*_-*comK, P*_*mtl*_-*comK*, and *P*_*hyp*_-*comK* constructs^8,9,13^. We also built *P*_*xyl47*_*-comK* and *P*_*T7*_-*comK* constructs to compare how high and low expression promoters impact competence induction. All constructs were integrated into *B. subtilis* along with the competence reporter (*P*_*comG*_-*YFP*) (**Figure 2a**). Adapting the induction protocol from Rahmer et. al^8^, we monitored competence reporter activity over time for each of the five strains at their respective maximum inducer concentrations (**Figure 2b**). Reporter activity begins to noticeably accumulate 1.5 h after induction which coincides with transforming DNA (tDNA) addition. YFP continues to accumulate throughout the duration of the experiment—though the rate of YFP production slows by the later time points. The reporter dynamics for each strain are similar; however, the *P*_*mtl*_-*comK* and *P*_*T7*_-*comK* strains show slightly slower induction of P_comG_ activity and lower maximum fluorescence. At lowered concentrations of inducer, the reporter demonstrates titratability in most strains **(Figure 2c**)^6^. However, the mannitol inducible strain shows remarkable sensitivity with 0.00019 % mannitol producing nearly the same response as 0.5 %. We also measured the reporter output produced when the Spizizen two-step competence protocol is utilized in the absence of inducible promoters (**Figure 2c**). At maximum output, fluorescence is 6.5× lower than that of *P*_*xyl*_-*comK* at 1% xylose. Previous work has shown that the Spizizen method results in a mixed population of competent and non-competent cells^14^. We anticipated that by using an inducible promoter to control competency, the entire population could enter the competent state. When high concentrations of inducers are present, we show that close to 96 to 99% of the population has strong reporter activity with a unimodal distribution for each of the five strains (**Figure 2d–e**). Low concentration of inducers results in large YFP negative populations (11–59 %) as well as large populations showing intermediate amounts of reporter activity (**Figure 2d–e**). The dynamics of the inducible competence strains contrast with that of WT under Spizizen conditions, which produces a clear bimodal population distribution with lower maximum fluorescence (**Figure 2e**).

**Figure 2.**
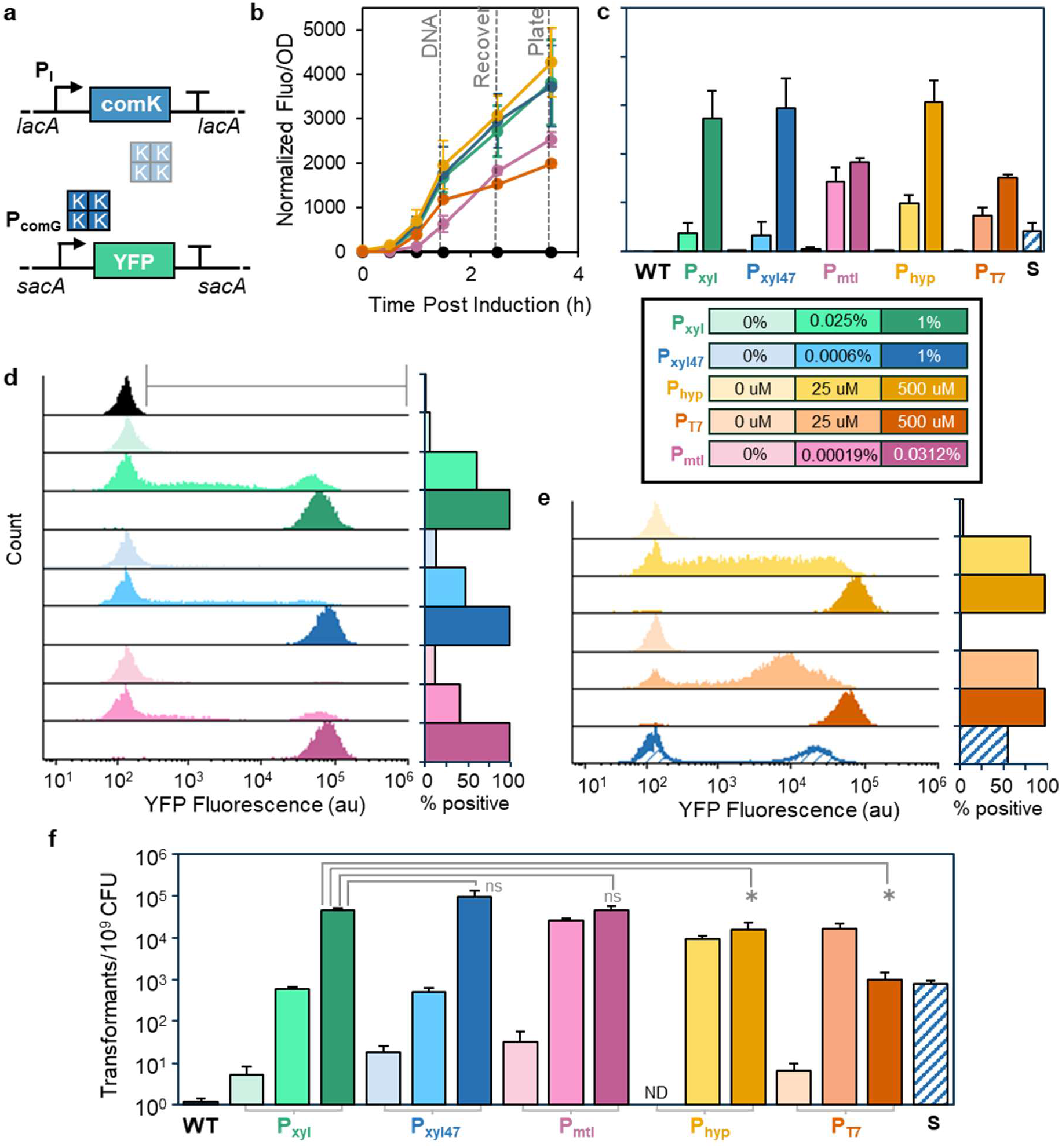
Comparison of common inducible promoters for competence induction. **a)** Construct design for inducible competence strains with competence reporter. **b)** Time course of OD_600_-normalized fluorescence of each strain at maximum inducer level. Induction occurs at 0 h, DNA is added at 1.5 h, fresh LB is added at 2.5 h, and cells are plated at 3.5 h. **c)** Reporter activity at 2.5 h obtained from the inducible promoters controlling ComK or WT under Spizizen protocol at equivalent time point. **d)** and **e)** Population distributions of reporter activity for each inducible condition or WT under Spizizen protocol at time of recovery. Counts are normalized to 100. Percent of the population displaying reporter activity defined by gray marker shown in (d). **f)** Transformation frequencies obtained from five inducible promoters controlling ComK expression. Statistical significance was determined by two-sided T-test (α = 0.05).

Finally, we directly compared the transformation frequencies of each strain under three inducer concentrations (**Figure 2f**). As suggested by the reporter data, very low concentrations of inducers can be sufficient to obtain medium levels of transformants. The activation at low concentration of inducer is likely due to interactions with the native *comK* locus where small amounts of induced ComK may be sufficient to trigger the positive feedback loop needed to enter the competent state^15^. The xylose and mannitol inducible strains have similar maximum transformation efficiencies— around 9 × 10^4^ transformants/10^9^ CFU; however, the two IPTG inducible strains were less efficient. The maximum efficiency for either strain is almost ten-fold lower. The differences between the xylose/mannitol and IPTG inducible strain efficiencies may be due to a reduction in total recoverable CFU (**Figure S1**). The Spizizen method resulted 56× lower transformation frequency than *P*_*xyl*_-*comK* at 1% xylose **(Figure 2f**). Together with the flow cytometric analysis, this suggests that the inducible promoters succeed at shifting most of the population into the competent state and, therefore, increase transformation frequency.

### Early ComK induction negatively impact transformation frequency

To further understand the induction dynamics of the high frequency inducible competence strains, we characterized the *P*_*xyl*_-*comK* and *P*_*mtl*_-*comK* strains—intending to capture any differences between the two sugar inducers. The dose responses for both promoters have surprising dynamics: the highest number of transformants are obtained at low concentrations of inducer—0.0625 % for P_xyl_ and 0.00019 % for P_mtl_ (**Figure 3a–b**). Interestingly, the number of transformants is commensurate with the total number of viable CFU recovered. At high concentration of inducers, growth rates are reduced, suggesting a high proportion of the population has entered the growth-arrested competent state (**Figure 3c–d**). In this instance, low growth rates appear to be limiting the total CFU recovered.

**Figure 3.**
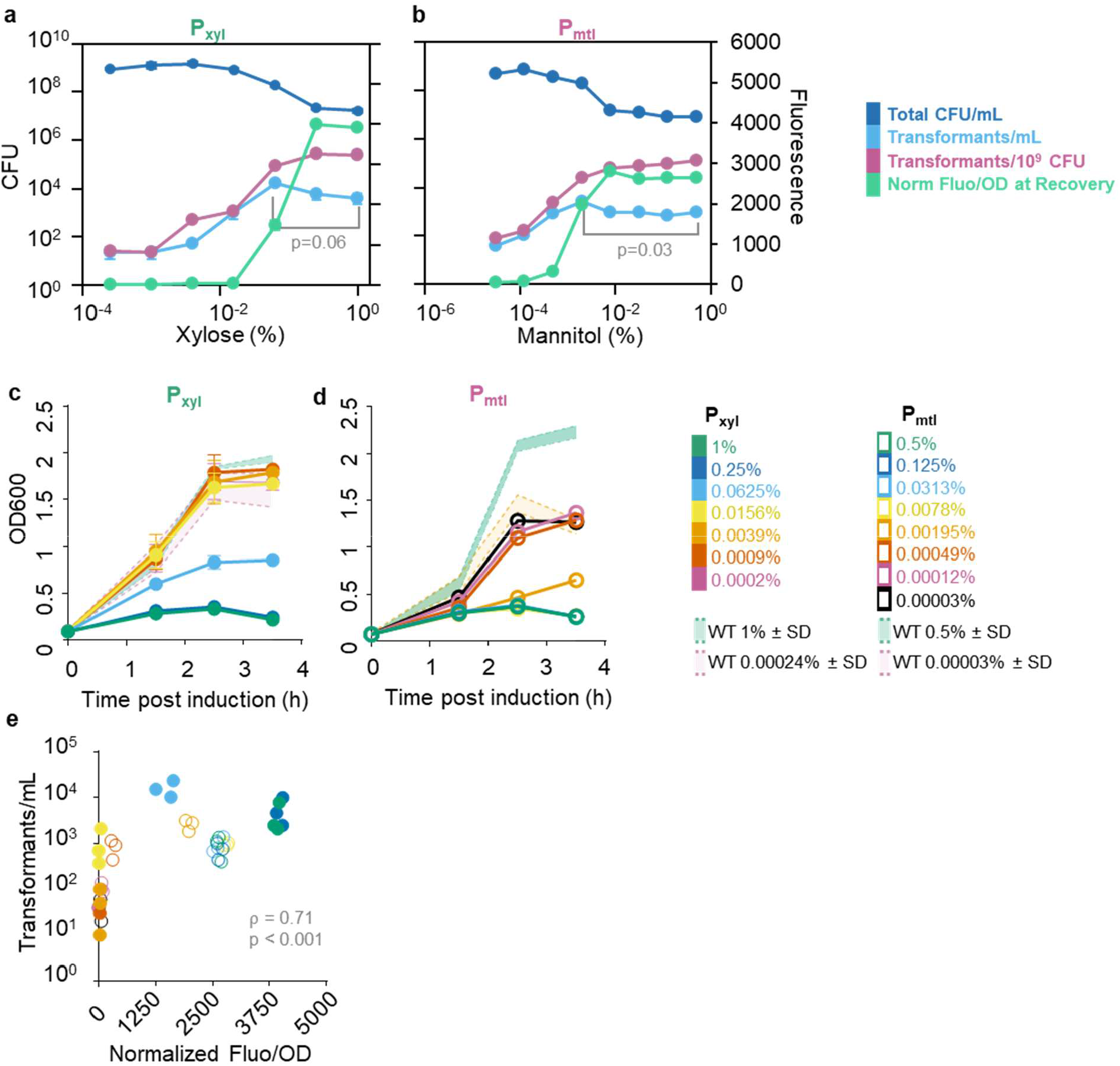
Trade-off between induction and total number of CFU driving transformation efficiency. **a)** Dose response curves for *P*_*xyl*_*-comK* and **b)** *P*_*mtl*_*-comK* strains. Statistical significance was determined by two-sided T-test (α = 0.05). **c)** OD_600_ growth curves of *P*_*xyl*_*-comK* and **d)** *P*_*mtl*_*-comK* strain at various concentrations of inducer. **e)** Correlation between transformants/mL and reporter activity at 2.5 h. Spearman’s rank correlation used to test the relationship.

The correlation between number of transformants and reporter activity at 2.5 h post-induction (time of recovery) is non-linear but somewhat monotonic (Pearson’s ρ = 0.71, p < 0.001) (**Figure 3e**). Extending recovery time demonstrates how cell growth affects the number of transformants obtained (**Figure S2**). At higher inducer concentrations, it takes at least 4 h to exit the competent state and begin growing. However, once the cells begin to divide, the transformation frequency improves in the *P*_*mtl*_-*comK* strain when induced at 0.03125 %. The *P*_*xyl*_-*comK* strain does not demonstrate an improvement in transformation frequency at longer recovery lengths; this may be due to differences in inducer persistence over time.

### Protocol optimization significantly increases transformation frequency

Given the apparent trade-off between maximum induction and sufficient growth in obtaining high numbers of transformants, we hypothesized that higher efficiencies could be obtained by increasing cell density prior to induction. To assess this, we grew the *P*_*xyl*_-*comK* strain for 2 h, as before, but concentrated at time of induction to various densities (**Figure 4a**). Concentration up to 9-fold increased the number of transformants 7-fold; however, the transformation frequency dropped as we increased cell density further. To streamline the protocol, we attempted to increase cell density by extending the pre-culture length, showing greater improvement than cell concentration (**Figure 4b**). Extending the length of pre-culture from 2 to 3 h, showed remarkable improvements in both number of transformants (64×) and transformation frequency (9×) (**Figure 4b**). Pre-culture lengths longer than 3 h reduce frequency, indicating the inducible competence protocol is potentially sensitive to changes in growth phase. We further optimize our protocol by testing induction and recovery lengths—observing modest improvement by extending induction length but no change with longer recovery (**Figure 4c–d**). With the optimized protocol, the dose response curve for the *P*_*xyl*_-*comK* strain indicates maximum frequency is obtained at xylose concentrations higher than 0.25 % (**Figure 4e**). Even at high concentration of xylose, the total CFU does not drop as much as seen in **Figure 3a**. The correlation between transformation frequency and reporter activity at three hours post induction is monotonic and non-linear, likely due to the positive feedback controlling expression from native *comK* locus (**Figure 4f**). Interestingly, the shift in protocol also changed the relationship between transformation frequency and reporter activity. With the optimized protocol, the reporter output is lower but associated with higher transformation frequency. Flow cytometric analysis of reporter activity shows similar behavior—xylose concentrations as low as 0.0625 % induces 98 % of the population into the competent state (**Figure 4g**). The protocol optimization described here improved number of transformants by 45× and transformation frequency by 19× compared to the original protocol induced with 0.0625% xylose (**Figure 4h**).

**Figure 4.**
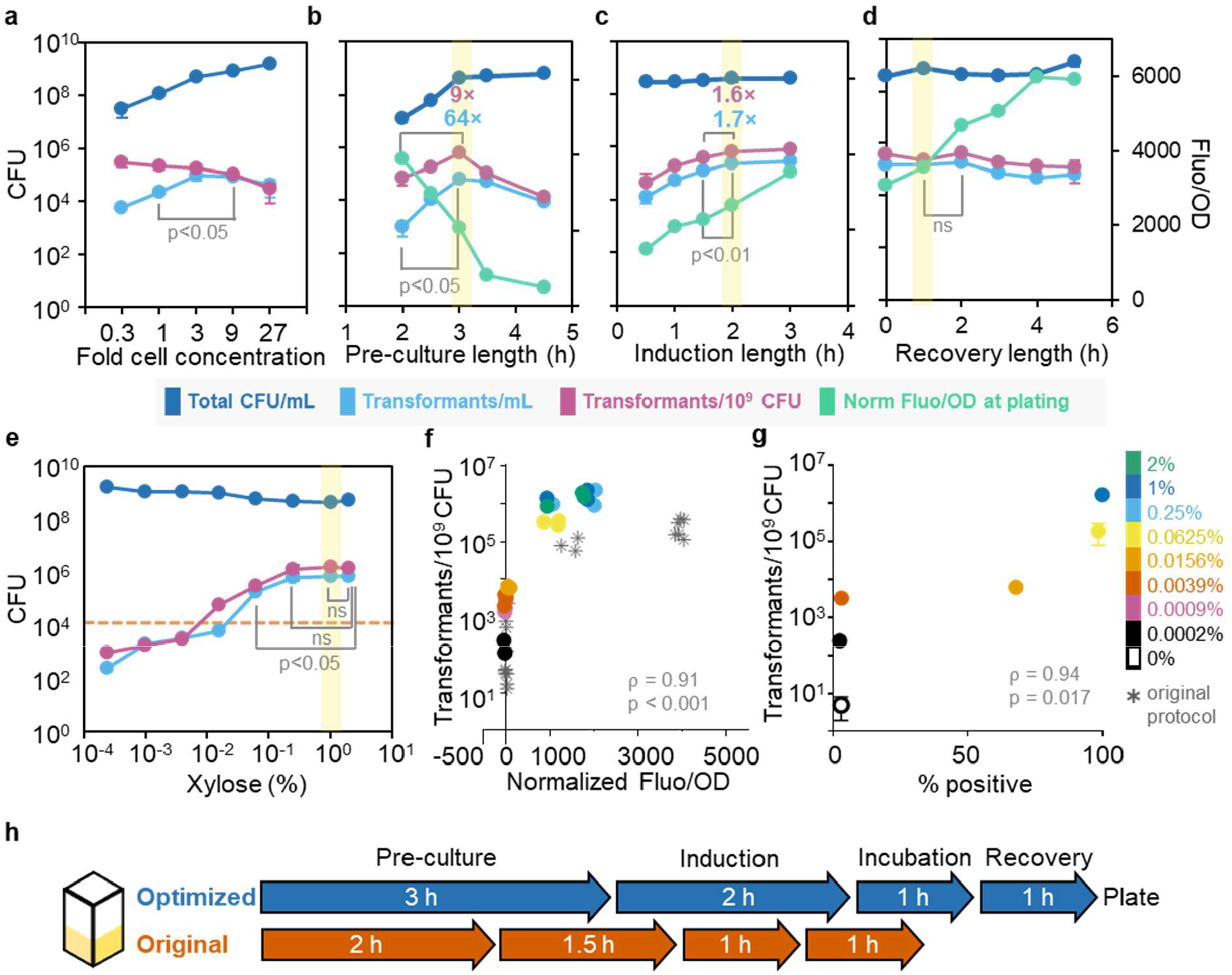
Optimized competence protocol improves transformation capacity and simplifies relationship with competence reporter. **a)** Effect of cell concentration on transformation in *P*_*xyl*_*-comK* strain after 2 h of pre-culture. **b)** Transformation efficiencies resulting from different pre-culture lengths. Highlighted conditions are carried over to subsequent figures. **c)** Effect of induction length on transformation after 3 h of pre-culture. **d)** Effect of recovery length on transformation after 3 h of pre-culture and 2 h induction duration. **e)** Dose response curve for *P*_*xyl*_*-comK* under optimized protocol conditions. The orange line corresponds to transformants/mL of *P*_*xyl*_*-comK* under the original protocol with 0.0625% xylose. **f)** Correlation between transformation frequency and normalized fluorescence per OD_600_ at time of recovery for the original and optimized protocols. **g)** Correlation between transformation frequency and percent of the population with competence reporter activity 3 h after induction. **h)** Schematic comparison of the original and optimized protocol step lengths.

### Disruption of native competence regulation reduces transformation frequency

ComK is transcriptionally, post-transcriptionally, and post-translationally regulated by several proteins^16,17^. Regulators of ComK are necessary to coordinate cell fate decision making under stressful conditions and to keep the number of ComK molecules low in the cell during exponential growth^7^. Several groups have knocked-out ComK negative regulators and seen changes to ComK expression or improvements in transformation frequency with the Spizizen method and inducible strains^8,14,17,18^. We chose to delete *rok, kre, codY*, and *degU* to test if the inducible competence strains could benefit from relieved repression of ComK (**Figure 5a**). *degU* was of special interest since Rahmer et al. saw a 4-fold improvement in a *P*_*mtl*_-*comKS* strain when *degU* was knocked-out^8^.

**Figure 5.**
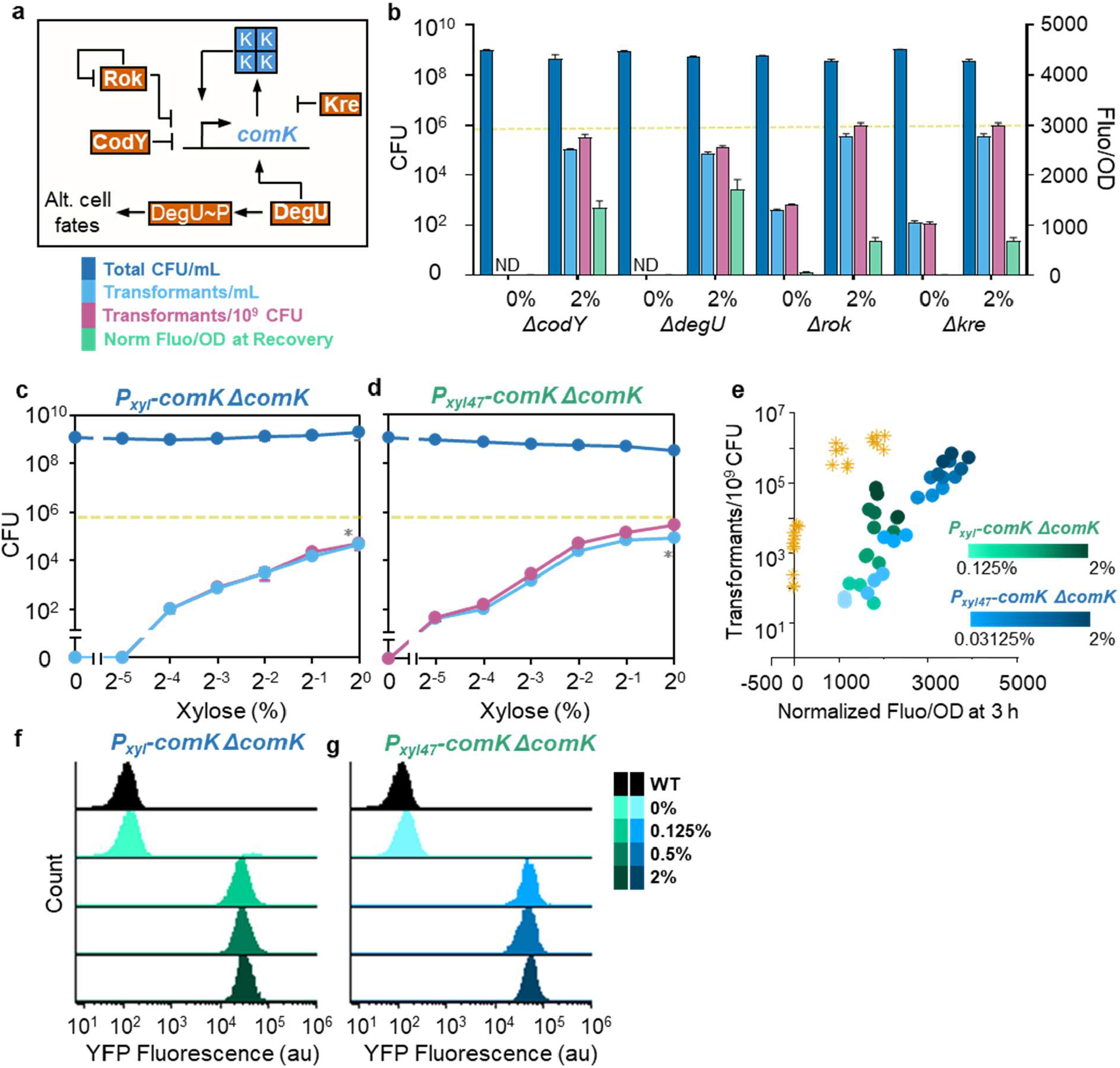
Disruption of native competence regulation does not improve transformation efficiency and shifts relationship with the competence reporter. **a)** A schematic depicting how several negative regulators impact ComK expression. **b)** Effect of negative regulator knockouts on transformation and reporter activity at 3 h. The yellow line indicates transformants/mL obtained with *P*_*xyl*_*-comK* under the optimized protocol at 2 %. **c)** Dose response characterization of *P*_*xyl*_*-comK* and **d)** *P*_*xyl47*_*-comK* strains in native Δ*comK* background. The yellow line indicates transformants/mL obtained with *P*_*xyl*_*-comK* under the optimized protocol at 2%. * indicates p < 0.05 for a two-sided T-test comparing transformants/mL to the optimized protocol at 2 %. **e)** Correlation between transformation frequency and reporter activity at 3 h in different genetic backgrounds. The yellow start data series corresponds to P_*xyl*_*-comK* at various xylose concentrations under the optimized protocol. **f)** Population distribution of reporter activity 3 h post-induction for *P*_*xyl*_*-comK* and **g)** *P*_*xyl47*_*-comK* Δ*comK* strains.

Under our optimized protocol conditions, the four knockouts did not improve transformation frequency. Rather, we saw a decrease in the number of transformants in Δ*codY* and Δ*degU* backgrounds with the *P*_*xyl*_-*comK* system (**Figure 5b**). We then tested if the P_xyl_ promoter provided sufficient ComK to overcome dependence on the native *comK* locus through knockout. The knockout resulted in a 15-fold decrease in the number transformants obtained (**Figure 5c**). Since we noticed improvements in transformation from 1 to 2 % xylose, we hypothesized that ComK expression is limiting in the knockout background. Using the higher expression P_xyl47_, we re-ran the dose response, obtaining a higher number of transformants, but not achieving the same efficiency as before (**Figure 5d**). The *comK* knockout also shifts the relationship between transformation frequency and reporter activity. The knockout strains have much higher fluorescence but lower transformation frequencies (**Figure 5e**). At a population level, the reporter shows monotonic, highly fluorescent populations—even at lower levels of inducer (**Figure 5f–g**).

### Higher transformation frequency enables construction of dual protein spore display library

Co-transformation of selection marker-less constructs allows for rapid multiplex genome editing; however, previous work in *B. subtilis* has been limited by low transformation (3 in 1 million) and co-transformation frequencies (2–5 %)^19^. To assess if the xylose inducible competence strain could perform better, we co-transformed spore protein (CotG and CotY) GUS fusion constructs with 2 kb homology arms for allelic replacement at their native loci. High transformation efficiencies are maintained in the presence of co-transforming DNA (ctDNA) (**Figure 6a**). Though frequencies varied slightly between the two loci, we found that 1 μg was sufficient ctDNA to obtain maximum co-transformation frequencies between 11–18% (**Figure 6b**). The inducible competence strain produces a greater number of co-transformants by creating a larger pool of transformants and more frequently integrating ctDNA. Leveraging the greater co-transformation frequency achieved through strain engineering and protocol optimization, we constructed and screened a spore display library of dual fluorescent proteins. Previous work has shown that the carrier/fusion partner, integration locus, and genetic background affects recombinant protein display levels on spores^19^. Furthermore, the design principles of displaying two different proteins—which would be useful for multi-enzyme pathway engineering—have yet to be characterized. To this end, we designed constructs for five spore proteins (CgeA, CotG, CotX, CotY, and SscA) to be fused to meYFP (YFP) and mCherry for integration at both their native loci and two orthogonal loci (*gltA* and *pyrD*) (**Figure 6c**). This library has 340 possible combinations of two integrations and 2760 possible combinations of three integrations. Our previous characterization of these five spore proteins for enzyme display showed SscA conferred the highest enzyme activity for soluble substrates^19^. However, SscA performed comparatively poorly on large, insoluble substrates due to low surface accessibility. Thus, SscA may not be appropriate for all applications. We decided to co-transform two subsets of spore fusion constructs: Library-24 (L24) without SscA and Library-30 (L30) with SscA constructs.

**Figure 6.**
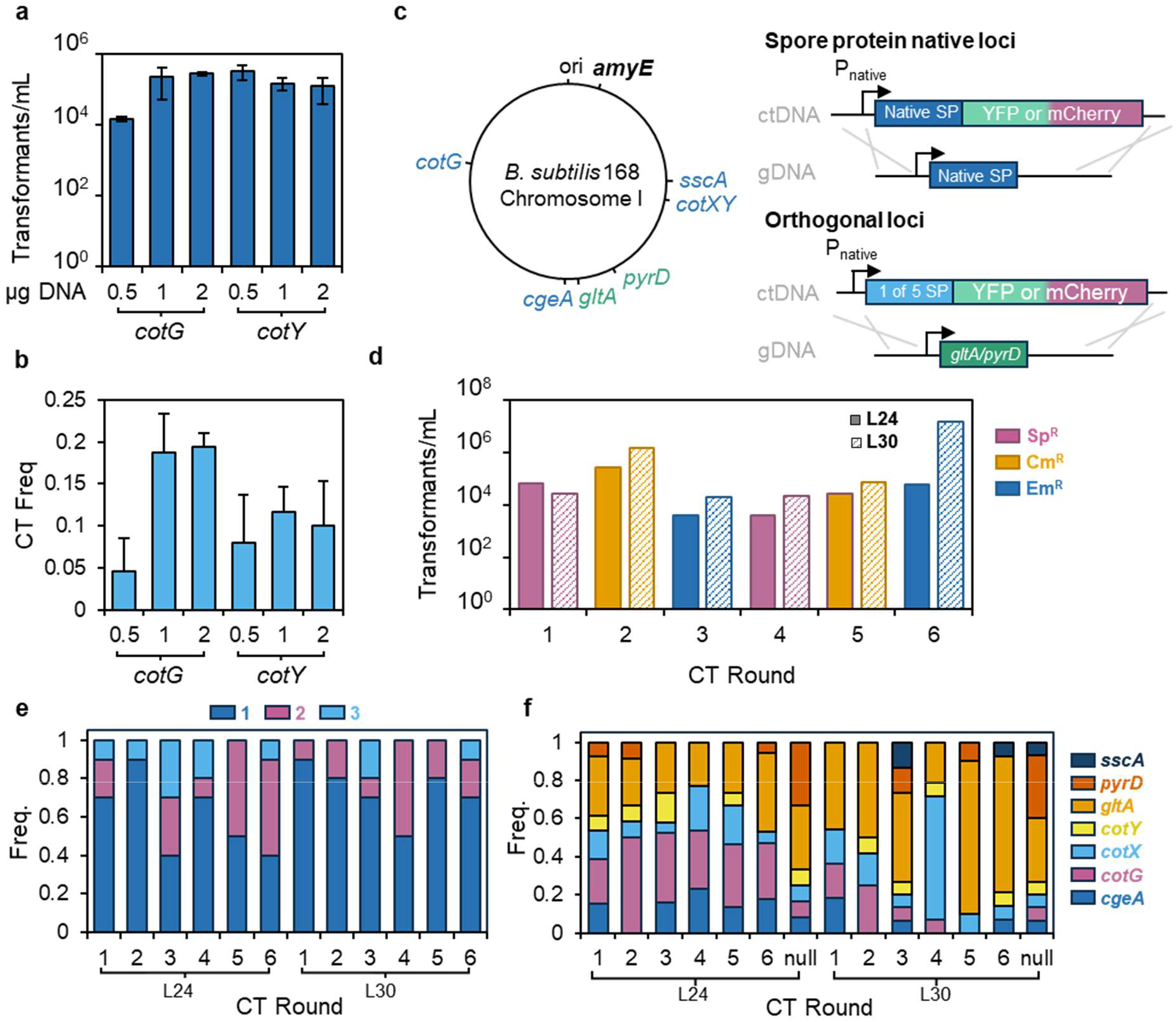
Dual protein spore display library design and construction. **a)** Transformants and **b)** co-transformation frequency obtained when pBS1C is co-transformed with variable amount of co-transforming DNA of CotG-GUS or CotY-GUS for allelic replacement. Co-transformants were quantified by blue-white screen on SG plates with X-Gluc. **c)** Chromosomal integration sites and construct design for spore protein-fluorescent protein fusions. Blue text indicates native spore protein loci and green text indicates orthogonal loci. **d)** Transformants/mL obtained over six rounds of co-transformation of Library-24 and Library-30. Sp = spectinomycin; Cm = chloramphenicol; Em = erythromycin. **e)** Frequency of integrations identified by MASC PCR over six rounds of co-transformation (n = 10). **f)** Frequency of integration at different loci over six rounds of co-transformation (n = 10). CT = co-transformation.

Starting from a pooled population of all single integrants, we conducted six rounds of co-transformation for each library, alternating between spectinomycin (Sp), chloramphenicol (Cm), and erythromycin (Em) resistances at *amyE*. We found that transformation of linearized plasmid DNA resulted in high frequencies of multi-resistance colonies; therefore, we transformed PCR products with 2 kb homology arms to minimize this occurrence. Over the six rounds of co-transformation, L24 showed very few instances of multi-resistant colonies, but L30 developed a large population of Sp^R^ + Em^R^ colonies during round 4 (89%) (**Figure S3**) that carried forward to rounds 5 and 6. The persistence of multi-antibiotic resistant colonies through rounds 4–6 explains abnormally high apparent transformation frequency observed for L30 in round 6 (**Figure 6d**). After each round, we measured the number of integrations by selecting random colonies for multiplex allele-specific combinatorial (MASC) PCR. We see the largest populations of multi-integrants after round 3 (**Figure 6e**). Subsequently, we see the measured co-transformation is slightly variable which may be due to sampling bias. Surprisingly, it was not rare to find triple integrations, even after only one round. For L24, we achieved a cumulative co-transformation frequency of 60 % after six rounds of co-transformation. All loci were transformable; however, the frequencies of integration were slightly different depending on locus (**Figure 6f**). For L24, integrations at *cotG* and *gltA* were overrepresented, whereas *pyrD* was underrepresented. For L30, *gltA* was also overrepresented, especially after round 4, which we attribute to an enrichment of multi-resistant colonies. To avoid overrepresentation of these multi-resistant clones, we decided to discard rounds 4–6 and use round 3 transformants for further analysis.

Selecting 186 random colonies from L24 round 6 and L30 round 3, we measured YFP and mCherry fluorescence directly from sporulated cultures (**Figure 7a,b**). We then selected 28 candidates based on the criteria of high YFP fluorescence, high mCherry fluorescence, or a combination of both for allelotyping by MASC PCR. We removed redundant strains and single integrations and then lysozyme purified the remaining strains to ensure that the signal was only from spores (**Figure 7c,d**). Compared to the fluorescence values of unpurified spores, the lysozyme purified strains had lower fluorescence per OD_600_ indicating much of the spore-fluorescent protein fusion may remain in mother cells, free floating in the media, or was only loosely associated with the spores (**Figure S4**). Nevertheless, the correlation between fluorescence of unpurified and lysozyme-purified fluorescence was high (Pearson’s r = 0.88 and 0.87 for YFP and mCherry). Among single integrants in L30, SscA promoted amongst the highest fluorescence at both its native locus and *gltA* (**Figure S5**). Other configurations that consistently promoted equivalent or higher fluorescence included *cotY-YFP/mCherry* and *cotY-YFP/mCherry* fusions at native and orthogonal loci. When considering concurrent display of both YFP and mCherry in this library, none of the multi-integrants were able to reach the same YFP fluorescence level as SscA was able to display individually. Nevertheless, the best combinations for dual display were combinations of SscA and either or both of CotG and CotX such as in L30-8 and L30-19 (**Figure 7c**).

**Figure 7.**
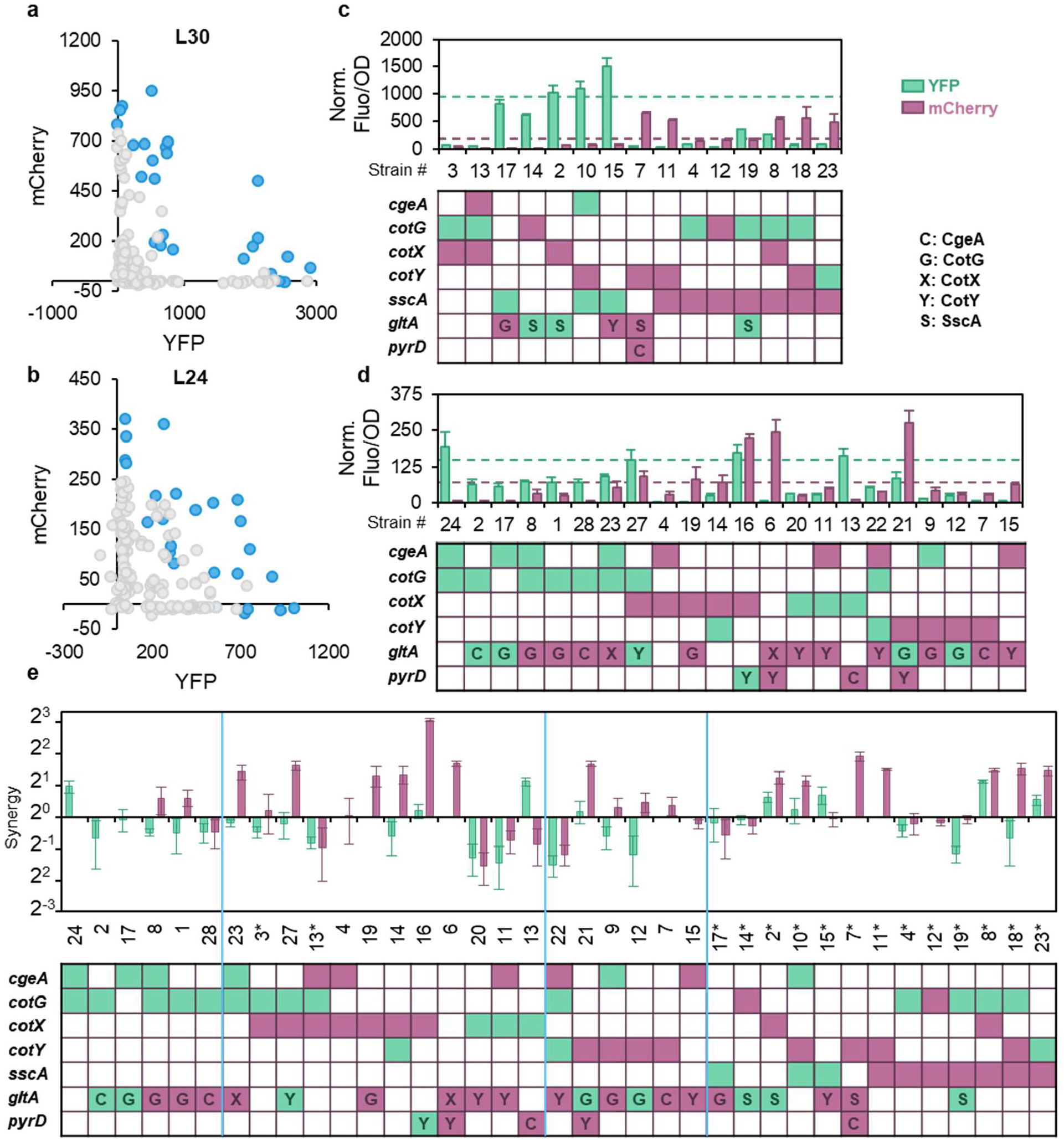
Analysis of top variants from co-transformation library screen. **a)** mCherry and YFP fluorescence from the screen of 186 random colonies from Library 30 and **b)** Library 24. Top variants are depicted in blue. **c)** Fluorescence of purified variants from Library 30 and **d)** Library 24. The green and purple lines represent CotY-YFP and mCherry (L30) or SscA-YFP and mCherry (L24), respectively. The table depicts the genotype of each strain. Green indicates YFP integration while purple indicates mCherry integration. The letters depict which spore protein was integrated at orthogonal loci. **e)** Fluorescence synergy of selected variants over the additive fluorescence of respective control strains. Variants from L30 are noted by (*).

Since SscA is not suitable for all applications, L24 provides alternative configurations for display of two proteins. Overall, fluorescence levels in this library were significantly lower than that for L30. Generally, CotG, CotX, and CotY are the best for YFP display, with the latter two being the best for mCherry display, as well. More so than in L30, L24 isolated strains show display is highly dependent on anchor protein, locus, and genetic background. L24-24 has the highest YFP fluorescence with *cotG::cotG-YFP* and *cgeA::cgeA-YFP* integrated; however, if CotG-YFP is integrated at *gltA* or mCherry fusions are integrated elsewhere, then YFP fluorescence decreases. L24-6 has the highest mCherry fluorescence with *gltA::cotX-mCherry* and *pyrD::cotY-mCherry* integrated. Combinations of CotX and CotY fusions work well to promote high fluorescence of both cargo proteins. The presence of CotY fusions at orthogonal loci especially seems to increase the display of CotX-mCherry. Though *pyrD* was underrepresented in the random MASC screen, it facilitates high fluorescence CotY fusions. L24-16 and L24-27—both CotX/CotY fusion containing strains—are the best performing dual fluorescence isolates in this library. While L24-16 (*cotX::cotX-mCherry, pyrD::cotY-YFP*) has high dual display, L24-11 and −20 perform poorly, highlighting anchor/cargo protein pairing and locus-dependent effects. Both L24-11 and 20 have CotX-YFP and CotY-mCherry integrated which severely lowers both display levels. Through analysis of L24 and L30, we show that variable amounts of display of two proteins can be achieved by selection of spore protein anchors and integration site.

Some combinations of spore fusions synergistically facilitated display of mCherry and YFP but most do not (**Figure 7e**). YFP fusions often enhance the display of mCherry fusions, to their own detriment. Strains containing *sscA-YFP/mCherry* fusions generally displayed high amount of synergy. Only three strains (L30-2, L30-8, and L30-23) synergistically display both YFP and mCherry: all three contain SscA fusions, whereas no constructs from L24 showed appreciable synergy for both fluorescent proteins. Several complex factors like genetic background, spore protein-fusion abundance, loading capacity, and protein-protein interactions likely drive differences in display levels.

## DISCUSSION

Engineering in *B. subtilis* is limited by its relatively low transformation efficiency and multiplex genome engineering capabilities. Efforts to increase transformation efficiency in *B. subtilis* have centered around use of inducible promoters to control expression of ComK; however, limited data shows direct comparison of these different systems. Here, we test five inducible promoters showing that all can induce competence, but mannitol and xylose inducible promoters are nearly 10-fold more efficient than the IPTG promoters. The IPTG inducible strains have lower recoverable total CFU at maximum induction—the induction dynamics of these strains could result in greater growth stalling and thus fewer transformants. Furthermore, IPTG is the only inducer that is not natively metabolized by *B. subtilis*. Longer persistence of the inducer may interfere with exit from competency resulting in low cell viability and fewer transformants^13,20^. The xylose and mannitol inducible strains are best for obtaining maximum number of transformants while the IPTG inducible strains may be best for experiments requiring long-term induction. Very low inducer concentrations are required to obtain high levels of transformation. We posit that the positive feedback loop at the native *comK* locus is the cause of this behavior and very little additional ComK is required to trigger the feedback loop.

As anticipated, we find that use of inducible promoters with high levels of inducer results in nearly 100 % of the population displaying reporter activity. Commonly, the P_comG_ reporter is used as a proxy indicator for cells being in the competent state; however, the exact threshold at which any given cell becomes transformable is unresolved^13,14,21,22^. Our flow cytometric analysis suggests that using inducible promoters shifts nearly the entire population into the competent state, but more conclusive evidence could include binning and sorting subpopulations to determine transformability of the population distribution. If the entire population is transformable, questions arise as to why only 1 in 1,000 cells successfully integrate and display antibiotic resistance. Higher amounts of tDNA do not change transformation frequency and it is therefore not likely the limiting factor (**Figure S6**). A transformation frequency of 1 in 1,000 is comparable to that obtained in *Vibrio cholerae*.; however, there are reports of much higher co-transformation frequencies (>60 %) in *V. cholerae*^2^. There are also other species of naturally competent bacteria with higher transformation frequencies. *Vibrio natriegens* and *Acinetobacter baylyi* have reports of frequencies around 1 in 1,000^22^. In *B. subtilis*, DNA binding, processing, stability in the cytosol, and recombination frequency are all events which may be limiting transformation frequency^2,10,23–26^. Interventions in these processes may increase transformation and co-transformation frequency further. Nonetheless, the use of inducible promoters increases reporter activity and the portion of the population displaying activity which correlates with 1700-fold higher transformation frequency.

A major finding of our work is the large influence of growth conditions on transformability and reporter output. Through protocol optimization, we saw a 45-fold improvement in number of transformants and 19-fold higher transformation frequency. As competency is a growth arrested state, timing induction with sufficient cell density is key. This improvement cannot entirely be explained by more total cells present; the optimized protocol only results in 3-fold more total CFU than the original protocol at 0.0625 % xylose. The later induction coincides with greater density as the cells approach stationary phase (**Figure S7**). Quorum sensing pheromones like ComX, CSF, and Phr peptides accumulate at higher densities and have several regulatory inputs which relieve of repression on ComK^27,28^. Coordinating induction of ComK with quorum sensing inputs may explain the outsized impact of increasing pre-culture length.

With the modified protocol, we were surprised to see the relationship between reporter output and transformation frequency change. At time of recovery, the original protocol resulted in higher fluorescence associated with lower transformation frequency; the optimized protocol tightened the range of fluorescence values but increased the maximum transformation frequency. The optimized protocol has slower rates of fluorescence production compared to the original protocol (**Figure S8**); however, the reporter output is equivalent at time of plating. The lower reporter output could be explained by the slower growth rate observed with the original protocol (**Figure S8**). At higher growth rates, more protein dilution occurs; once the cells slow in growth post-3 h, there are equivalent amounts of fluorescence in both conditions^29^.

Previous studies have suggested expression of ComK from recombinant promoters is insufficient to overcome repression from negative regulators like *degU* or deletion of the native *comK* gene^8^. In our study, we find our strain does not benefit from deletion of negative regulators. Removing *rok* and *kre* did not have an impact on transformation frequency. Under Spizizen culture conditions, the *kre* deletion has been shown to have a 30-fold improvement in transformation frequency compared to WT^14^. However, the P_xyl_ promoter may be producing sufficient ComK levels to overcome the effect of negative regulators. The other two deletions, *codY* and *degU*, had detrimental impacts on transformation. Previous work had shown that deletion of *degU* can increase transformation frequency in inducible competence strains; however, the benefit/detriment of *degU* seems to be context dependent^8^. DegU has multiple roles during cell fate decision making. Unphosphorylated/low-levels of phosphorylated DegU can promote ComK expression by facilitating ComK binding to its own promoter^30^. Alternatively, high levels of phosphorylated DegU increases exoprotease expression and promotes biofilm formation^23^. Under the growth conditions used in this study, DegU could be assisting early expressing ComK with binding to the native promoter. Thus, DegU is required to achieve the maximum number of transformants. Neither P_xyl_ nor P_xyl47_-*comK* were sufficient to overcome the deletion of the native *comK* locus, though P_xyl47_-*comK* was better performing in the Δ*comK* background. The suboptimal performance of the inducible strains could be due to insufficient ComK expression from the inducible promoters alone or shifting the period in which the population is most transformable. We tested combining the *comK* knockout with that of negative regulators and saw no improvement, indicating low expression is not likely the only limitation (**Figure S9**). Reporter activity throughout the experiment shows differences compared to when the *comK* locus is intact (**Figure S9**). The reporter turns on faster, behaving more similarly to the original protocol; however, the signal begins to be diluted out by time of plating. Without the positive feedback loop, the inducible promoter may not have enough sustained expression to maintain competency. The discrepancies in reporter behavior in different genetic backgrounds also highlights the complex relationship between P_comG_ activity and actual transformation frequency.

Improvements in transformation efficiency facilitate multiplex genome engineering and engineering complex phenotypes like spore display. Not only does the P_xyl_-*comK* strain have 2800 × greater transformants than WT with the Spizizen method, but it also has 2–9 × greater co-transformation frequency^19^. These improvements enabled the co-transformation of two spore display libraries with 30 genomic integration constructs at seven loci. After multiple rounds of co-transformation, we were able to reach cumulative co-transformation frequencies as high as 60 %. Allelotyping by MASC PCR of 56 strains identified by initial fluorescence screen revealed complex design considerations for display of two recombinant proteins. Our work confirms that SscA is an excellent spore display anchor protein with higher display capacity than other commonly used spore proteins. However, for dual protein display, we observed a general tradeoff of diminished YFP fluorescence when mCherry fusions are present. This may be due to limitations in expression capacity and competition during spore coat assembly. Even under these constraints, our screen identified several strains with high display of both YFP and mCherry in both libraries. Strains containing SscA fusions also support the greatest synergy in both YFP and mCherry display. Furthermore, the several co-display strains demonstrate different relative fluorescence indicating the results could inform design of future enzyme cascades on spores. Previous work has shown the important of optimizing enzyme stoichiometry for efficient biocatalyst cascades^32^.The display of enzyme cascades on spores is an ongoing effort, limited by poor characterization of how multi-fusion strains function. Our work shows that spore anchor, genetic background, integration locus, presence of additional spore coat protein copies, and cargo protein identity all influence display capacity. CotX and CotY are commonly utilized coat proteins for spore display due to their localization to the outermost crust layer^33^. But, CotX/Y/Z are key morphogenic proteins for crust formation^34,35^. While CotX/Y/Z may have some redundant functionality during crust formation, the loading of other crust proteins may be limited in the presence of any knockout^34^. We observe that CotY fusions benefit from integration at orthogonal loci, more so than CotX fusions. CotY may not be able to carry out its morphogenic functionalities when fused to recombinant proteins— ultimately limiting display capacity of the crust. This may be a contributing factor to why CotY fusions at *gltA* and *pyrD* result in higher display levels along with potential differences in gene expression. Understanding the underlying principles driving spore display capacity will be important to the development of more complex enzymatic cascades on spores. High throughput screening facilitated by high transformation efficiencies is a promising route to optimizing display of recombinant proteins on spores.

## MATERIALS AND METHODS

### Bacterial strains and culture conditions

*B. subtilis* strain 168 (Δ*trpC2*) was used to derive all strains described in this study. Strains, plasmids, and oligonucleotides used in this study are listen in **Tables S1–S3**. For plasmid construction and propagation, *E. coli* DH5α was used. Cells were grown in LB medium at 37 °C. Antibiotics were used at the following concentrations: chloramphenicol (Cm) 5 μg/mL, kanamycin 10 μg/mL, spectinomycin (Sp) 100 μg/mL, erythromycin (Em) 1 μg/mL, zeocin 100 μg/mL, and ampicillin 100 μg/mL.

### Transformation

Preparation of naturally competent *B. subtilis* cells was done as described previously^6^. Transforming DNA was prepared by one of two methods. 1) Plasmid DNA was miniprepped from *E. coli* and linearized by restriction enzyme (either ScaI-HF or SacI). 2) Transformation constructs were assembled by overlap-extension PCR and purified by gel extraction. For both methods, 100 ng of tDNA was used. For inducible competence, overnights were set from freshly streaked colonies on LB containing appropriate antibiotics. After 24 h, cells were subcultured 1:25 into 3 mL fresh LB. The original transformation protocol is as follows: 2 h pre-culture with shaking, inducer added, 1.5 h induction duration, tDNA added to 500 μL of cells, 1 h incubation, 500 μL of LB added, 1 h recovery, dilute cells and plate. Cells were diluted to obtain 20–200 cells per plate. Transformation frequency was calculated based on the ratio of transformants to total CFU on antibiotic-free plates. The competence reporter was assessed by microplate reader (SpectraMax M3) measuring absorbance (600 nm) and fluorescence (ex. 485, em. 525 nm).

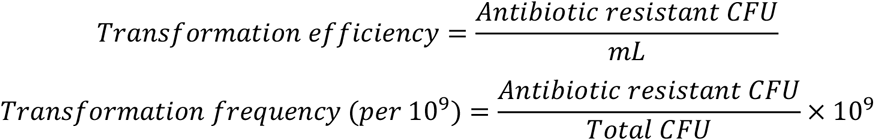

Co-transformation frequency of GUS fusion constructs was measured by co-transforming 100 ng linearized pBS1C with 0.5 to 2 μg of linearized pENTR *cotG-C-GUS* or *cotY-C-GUS* for integration at their respective native loci. Transformations were plated on 2× SG plates containing 100 μg/mL X-Gluc and number of blue and white colonies were counted after 48 h.

### Flow Cytometry

Flow cytometry was performed on an Attune NxT flow cytometer. At least 10,000 events were recorded per sample. A green laser (488 nm) at a voltage of 340 V was used to record fluorescence, and gating was performed on a FSC-H vs. SSC-H plot.

### Co-Transformation Library Preparation and Screening

DNA constructs for the co-transformation libraries were prepared as follows for all except those containing SscA: miniprep from *E. coli* DH5α, linearize with EcoNI or DrdI, concentrate with butanol extraction, and purify by ethanol precipitation. SscA containing plasmids were unstable in *E. coli*; therefore, those constructs were PCR amplified and purified by column purification. DNA was resuspended in TE buffer. Each co-transformation construct was pooled and 1 μg of each construct was co-transformed with 300 ng of PCR linearized chloramphenicol, spectinomycin, or erythromycin resistance marker with 2 kb homology arms for *amyE*. For each round of co-transformation, the xylose inducible protocol described above was utilized. After 1 h of recovery, cells were plated on the current antibiotic for transformation efficiency. The remaining transformation was subcultured 1:50 in fresh LB with antibiotic. After overnight growth, the outgrowth was plated on all antibiotics and either frozen or re-transformed. Random colonies were selected from LB antibiotic plates and placed in 24 deep well plates for growth shaking overnight at 37 °C. Overnight cultures were subcultured 1:50 into 2× DSM (16 g mL^-1^ Diffco Nutrient Broth, 0.2 % KCl, 0.024 % MgSO_4_·7H_2_O, 0.01 mM MnCl_2_·4H_2_O, 1 mM Ca(NO_3_)_2_·4H_2_O, 1 μM FeSO_4_·7H_2_O, pH adjusted to 7.6). After 24 hours shaking at 37 °C, spores were diluted 10× in DI water. OD_600_, mCherry fluorescence (ex. 585 nm, em. 615 nm), and YFP fluorescence (ex. 485 nm, em. 525 nm) were measured by microplate reader.

### MASC (multiplexed allele-specific combinatorial) PCR

To validate the MASC PCR primer pools, each individual primer pair was tested for specificity and sufficient efficiency on single integrant strains. All MASC reactions were done with OneTaq Quick Load in standard buffer. Single colonies were resuspended in 20 mM NaOH, held at 95 °C for 20 minutes, and diluted 1:50 in water. 20 μL Reactions contained 2 μM of each primer and 1 μL of diluted template. Touchdown cycling conditions reduces the amount of off target amplification and was as follows: 94 °C for 3:30 min, (94 °C for 30 s, 68 °C for 30 s, 68 °C for 2 min) × 5 cycles, (94 °C for 30 s, 64 °C for 30 s, 68 °C for 2 min) × 5 cycles, (94 °C for 30 s, 60 °C for 30 s, 68 °C for 2 min) × 20 cycles. Reactions were run on 1.5 % agarose gels containing SYBR Safe for visualization. The MASC PCR assay is composed of five reactions per colony: WT, *mCherry, YFP, gltA*, and *pyrD* (**Figure S10**). All primer pools were specific with few unintended bands; however, the *mCherry* and *YFP* on *cotX-mcCherry/YFP* templates (target amplicon size 600 bp) also produces a 3 kb amplicon due to the proximity with *cotY*. Additionally, the *pyrD* primers in the WT pool produces an unexpected 1.5 kb amplicon; however, this band does not overlap with any intended amplicons and does not interfere with the assay. The WT pool of primers contains forward primers for each of the seven loci with 3’ regions binding to the stop codon and a few downstream base pairs for native spore protein loci and 3’ regions binding to the start codon and a few downstream base pairs for orthogonal loci. The reverse primers of the WT, *mCherry*, and *YFP* pools are the same with each primer binding 200, 400, 800, 1000, 1200, or 2000 base pairs downstream of the forward primers. The forward primers for the *mCherry* and *YFP* pools are a single primer binding either *mCherry* or *YFP* coding sequence. For the *gltA* and *pyrD* pools, there are five forward primers specific to the coding sequence of each spore protein and one reverse primer specific to each locus. The intended amplicon sizes are 200, 400, 600, 800, and 1000 bp. The WT pool confirms which loci have been edited, the mCherry and YFP pools confirm which loci have which fluorescent proteins, and the *gltA* and *pyrD* pools confirm which spore proteins are integrated at the orthogonal loci.

### Spore Purification

Spores were grown as described above. To isolate spores from vegetative/mother cells, spores were centrifuged at 3,000 ×g for 5 min, washed once with DI water, and incubated with 5 μg/mL lysozyme in 50 mM Tris-HCl buffer (pH 7.2) for 15 min. Spores were then washed again with DI water and subsequently washed with a 0.05 % (w/v) SDS solution. After the SDS was removed, three DI water washes followed.

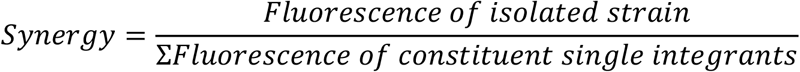

## AUTHOR CONTRIBUTIONS

Jessica A. Lee – Conceptualization; formal analysis; investigation; methodology; data visualization; writing; editing. Nikhil U. Nair – Conceptualization; funding acquisition; supervision; writing—review & editing.

## SUPPORTING INFORMATION

Supplemental Information: All supplemental figures and tables Supplementary Data 1: Sequence information of all integration cassettes.

## ACKNOWLDGEMENTS

We thank all past and present members of the Nair lab for their helpful discussion. We would also like to thank Prof. James A. Van Deventer (Tufts ChBE), Prof. Andrew Camilli (Tufts GSBS), and Prof. Lauren Andrews (UMass-Amherst) for their thoughtful feedback on experimental designs. This work was supported by CBET-2208390 (NSF), CBET-1935354 (NSF), BioMADE (T-ADP24-A-1166), and DARPA (contract HR00112590052) to N.U.N.

## REFERENCES

1. Lopez, D., Vlamakis, H. & Kolter, R. Generation of multiple cell types in Bacillus subtilis. FEMS Microbiol. Rev. 33, 152–163 (2009).

2. Dalia, A. B., McDonough, E. & Camilli, A. Multiplex genome editing by natural transformation. Proc. Natl. Acad. Sci. 111, 8937–8942 (2014).

3. Dalia, T. N. et al. Multiplex genome editing by natural transformation (MuGENT) for synthetic biology in Vibrio natriegens. ACS Synth. Biol. 6, 1650–1655 (2017).

5. Koo, B.-M. et al. Construction and Analysis of Two Genome-scale Deletion Libraries for Bacillus subtilis. Cell Syst. 4, 291–305.e7 (2017).

6. Spizizen, J. TRANSFORMATION OF BIOCHEMICALLY DEFICIENT STRAINS OF BACILLUS SUBTILIS BY DEOXYRIBONUCLEATE. Proc. Natl. Acad. Sci. 44, 1072–1078 (1958).

7. Maier, B. Competence and Transformation in *Bacillus subtilis*. Curr. Issues Mol. Biol. 57–76 (2020) doi:10.21775/cimb.037.057.

8. Rahmer, R., Morabbi Heravi, K. & Altenbuchner, J. Construction of a Super-Competent Bacillus subtilis 168 Using the PmtlA-comKS Inducible Cassette. Front. Microbiol. 6, 1431 (2015).

9. Zhang, X.-Z. & Zhang, Y.-H. P. Simple, fast and high-efficiency transformation system for directed evolution of cellulase in Bacillus subtilis. Microb. Biotechnol. 4, 98–105 (2011).

10. Deng, A. et al. Simultaneous Multiplex Genome Engineering via Accelerated Natural Transformation in Bacillus subtilis. Front. Microbiol. 12, (2021).

11. Castillo-Hair, S. M., Baerman, E. A., Fujita, M., Igoshin, O. A. & Tabor, J. J. Optogenetic control of Bacillus subtilis gene expression. Nat. Commun. 10, 3099 (2019).

12. Castillo-Hair, S. M., Fujita, M., Igoshin, O. A. & Tabor, J. J. An Engineered B. subtilis Inducible Promoter System with over 10 000-Fold Dynamic Range. ACS Synth. Biol. 8, 1673–1678 (2019).

13. Espinar, L., Dies, M., Çağatay, T., Süel, G. M. & Garcia-Ojalvo, J. Circuit-level input integration in bacterial gene regulation. Proc. Natl. Acad. Sci. 110, 7091–7096 (2013).

14. Gamba, P., Jonker, M. J. & Hamoen, L. W. A Novel Feedback Loop That Controls Bimodal Expression of Genetic Competence. PLOS Genet. 11, e1005047 (2015).

15. van Sinderen, D. et al. comK encodes the competence transcription factor, the key regulatory protein for competence development in Bacillus subtilis. Mol. Microbiol. 15, 455–462 (1995).

16. Hahn, J., Luttinger, A. & Dubnau, D. Regulatory inputs for the synthesis of ComK, the competence transcription factor of Bacillus subtilis. Mol. Microbiol. 21, 763–775 (1996).

17. Hoa, T. T., Tortosa, P., Albano, M. & Dubnau, D. Rok (YkuW) regulates genetic competence in Bacillus subtilis by directly repressing comK. Mol. Microbiol. 43, 15–26 (2002).

18. CodY is required for nutritional repression of Bacillus subtilis genetic competence. https://journals.asm.org/doi/epdf/10.1128/jb.178.20.5910-5915.1996 doi:10.1128/jb.178.20.5910-5915.1996.

19. Chappell, T. C. et al. Advancing protein display on bacterial spores through an extensive survey of coat components. 2024.11.22.624950 Preprint at 10.1101/2024.11.22.624950 (2024).

20. Wiesler, E. E. et al. ComK-induced cell death is reversed by upregulating the SigB or Spx pathway in Bacillus subtilis. Microbiol. Spectr. 13, e01612–25 (2025).

21. Smits, W. K. et al. Stripping Bacillus: ComK auto-stimulation is responsible for the bistable response in competence development. Mol. Microbiol. 56, 604–614 (2005).

22. Boonstra, M. et al. Analyses of competent and non-competent subpopulations of Bacillus subtilis reveal yhfW, yhxC and ncRNAs as novel players in competence. Environ. Microbiol. 22, 2312–2328 (2020).

23. Ellison, T. J. & Ellison, C. K. Improved DNA binding to a type IV minor pilin increases natural transformation. Nucleic Acids Res. 53, gkaf467 (2025).

24. Blokesch, M. & Schoolnik, G. K. The Extracellular Nuclease Dns and Its Role in Natural Transformation of *Vibrio cholerae*. J. Bacteriol. 190, 7232–7240 (2008).

25. Dalia, T. N. et al. Enhancing multiplex genome editing by natural transformation (MuGENT) via inactivation of ssDNA exonucleases. Nucleic Acids Res. 45, 7527–7537 (2017).

26. Wendt, K. E., Walker, P., Sengupta, A., Ungerer, J. & Pakrasi, H. B. Engineering Natural Competence into the Fast-Growing Cyanobacterium Synechococcus elongatus Strain UTEX 2973. Appl. Environ. Microbiol. 88, e01882–21.

27. Pottathil, M., Jung, A. & Lazazzera, B. A. CSF, a Species-Specific Extracellular Signaling Peptide for Communication among Strains of Bacillus subtilis and Bacillus mojavensis. J. Bacteriol. 190, 4095–4099 (2008).

28. Schneider, K. B., Palmer, T. M. & Grossman, A. D. Characterization of comQ and comX, Two Genes Required for Production of ComX Pheromone in Bacillus subtilis. J. Bacteriol. 184, 410–419 (2002).

29. Nordholt, N., van Heerden, J., Kort, R. & Bruggeman, F. J. Effects of growth rate and promoter activity on single-cell protein expression. Sci. Rep. 7, 6299 (2017).

30. Hamoen, L. W., Van Werkhoven, A. F., Venema, G. & Dubnau, D. The pleiotropic response regulator DegU functions as a priming protein in competence development in Bacillus subtilis. Proc. Natl. Acad. Sci. U. S. A. 97, 9246–9251 (2000).

31. Marlow, V. L. et al. Phosphorylated DegU Manipulates Cell Fate Differentiation in the Bacillus subtilis Biofilm. J. Bacteriol. 196, 16–27 (2014).

32. Chen, L., Mulchandani, A. & Ge, X. Spore-displayed enzyme cascade with tunable stoichiometry. Biotechnol. Prog. 33, 383–389 (2017).

33. A Distance-Weighted Interaction Map Reveals a Previously Uncharacterized Layer of the Bacillus subtilis Spore Coat. Curr. Biol. 20, 934–938 (2010).

34. Imamura, D., Kuwana, R., Takamatsu, H. & Watabe, K. Proteins Involved in Formation of the Outermost Layer of Bacillus subtilis Spores. J. Bacteriol. 193, 4075–4080 (2011).

35. Zhang, J., Fitz-James, P. C. & Aronson, A. I. Cloning and characterization of a cluster of genes encoding polypeptides present in the insoluble fraction of the spore coat of Bacillus subtilis. J. Bacteriol. 175, 3757–3766 (1993).

