## Supplemental Information for "Enhanced *Bacillus subtilis* natural competence enables multiplexed genome and spore engineering"

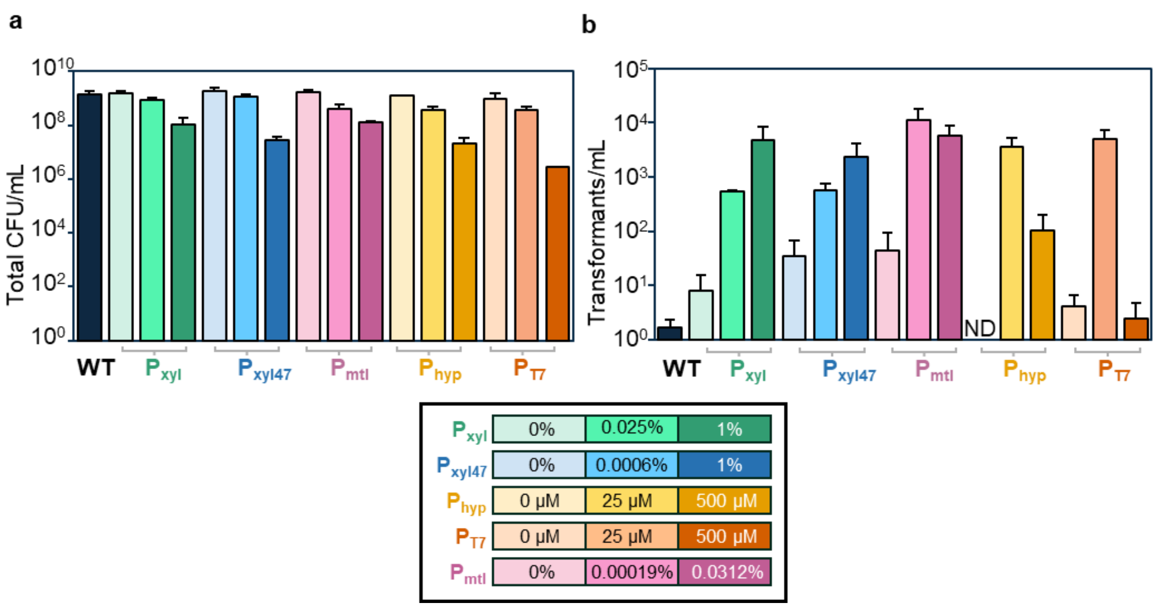


**Figure S1.** Comparison of inducible promoters for competence induction. a) Total CFU/mL recovered after transformations. b) Transformants/mL recovered after transformations.

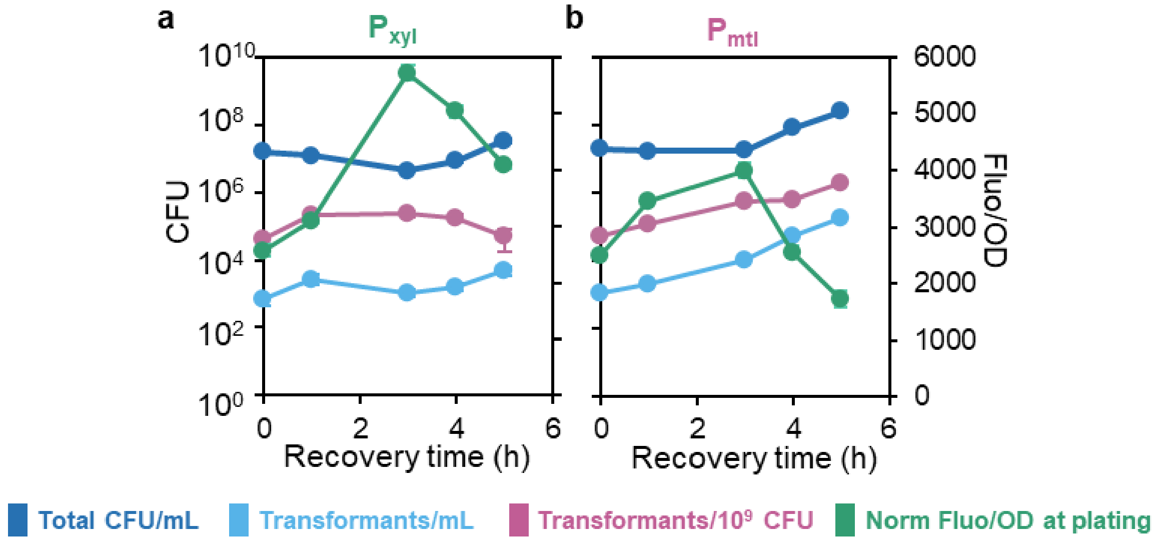


**Figure S2.** Effect of recovery time on transformation. a) $P_{xyl}$-*comK* with 1% xylose and b) $P_{mtl}$-*comK* with 0.03125% mannitol.

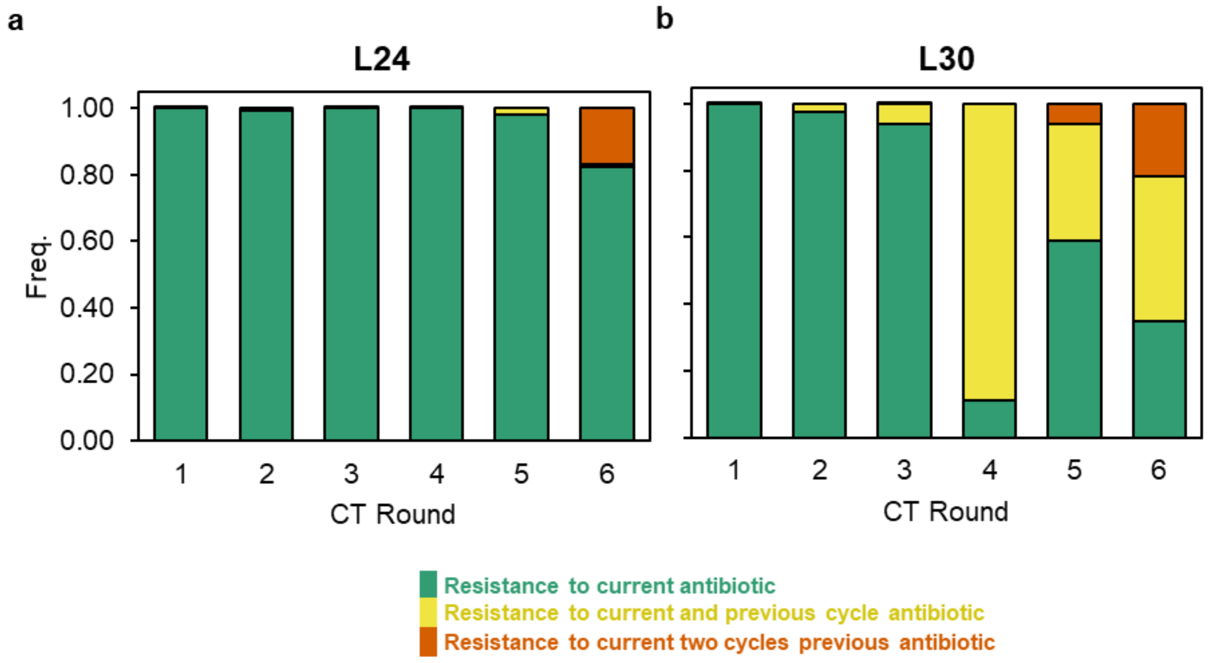


**Figure S3.** a) Effect of knockouts on transformation of $P_{xyl47}$-*comK* Δ*comK* strain. Dashed line represents transformants/mL of $P_{xyl47}$-*comK* Δ*comK* at 2% xylose. b) Normalized fluorescence per OD over time for $P_{xyl}$-*comK* Δ*comK*, and c) $P_{xyl47}$-*comK* Δ*comK*.

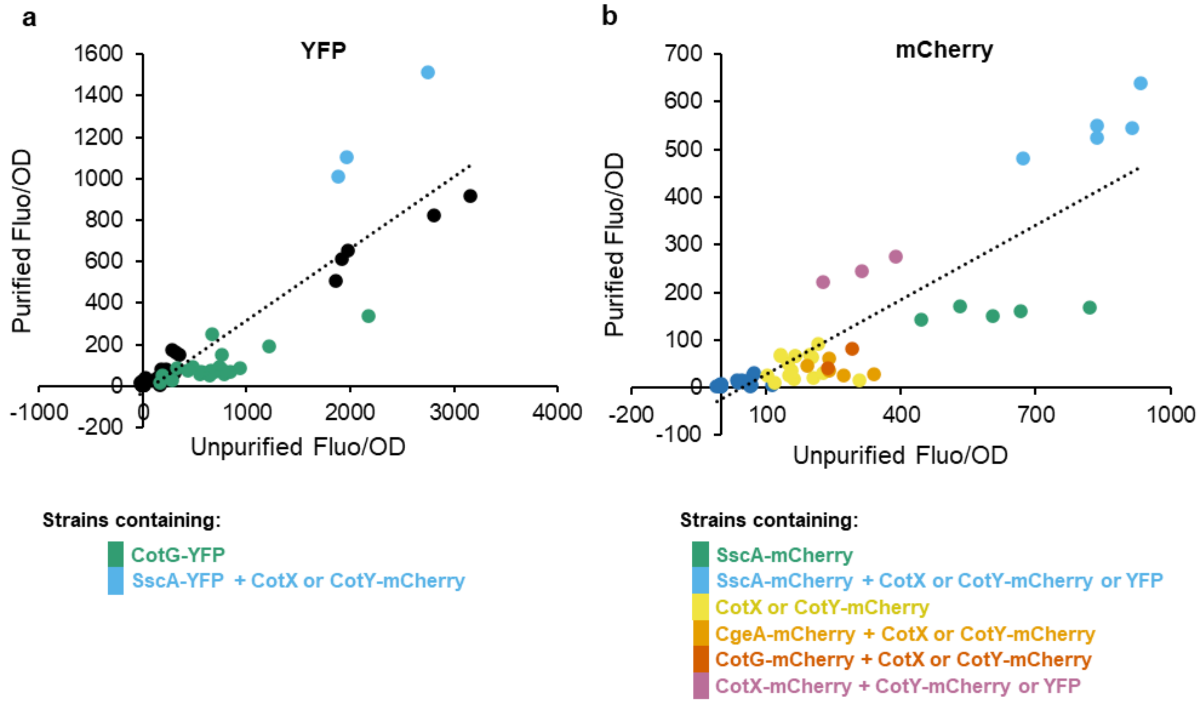


**Figure S4.** a) YFP and b) mCherry fluorescence of unpurified vs lysozyme purified spores.

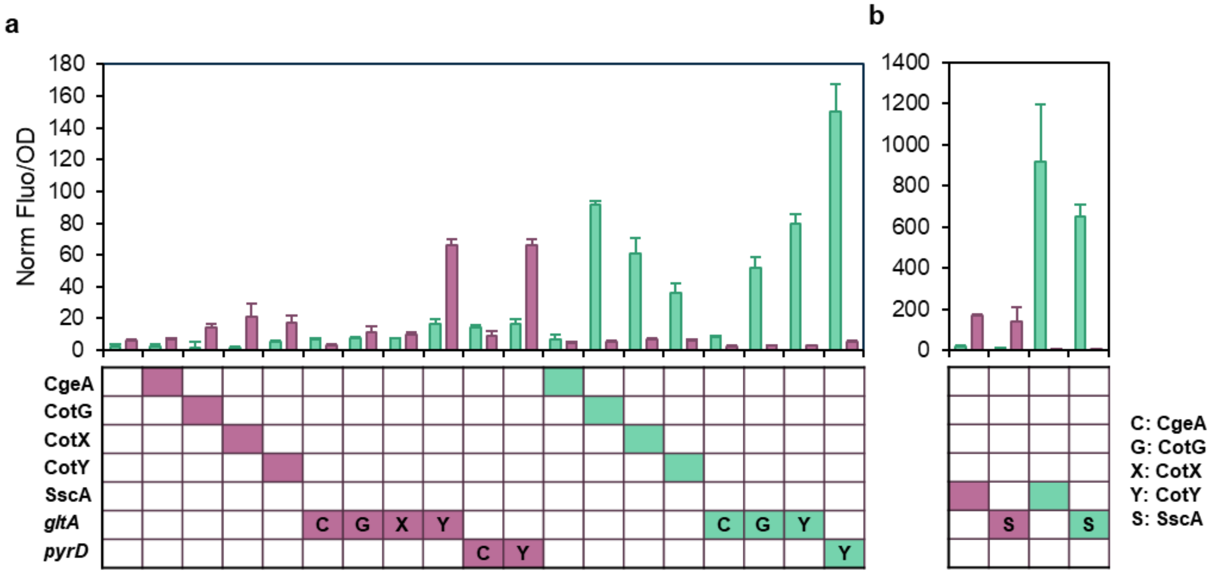


**Figure S5.** Single integration strains after lysozyme purification from a) Library 24 and b) Library 30.

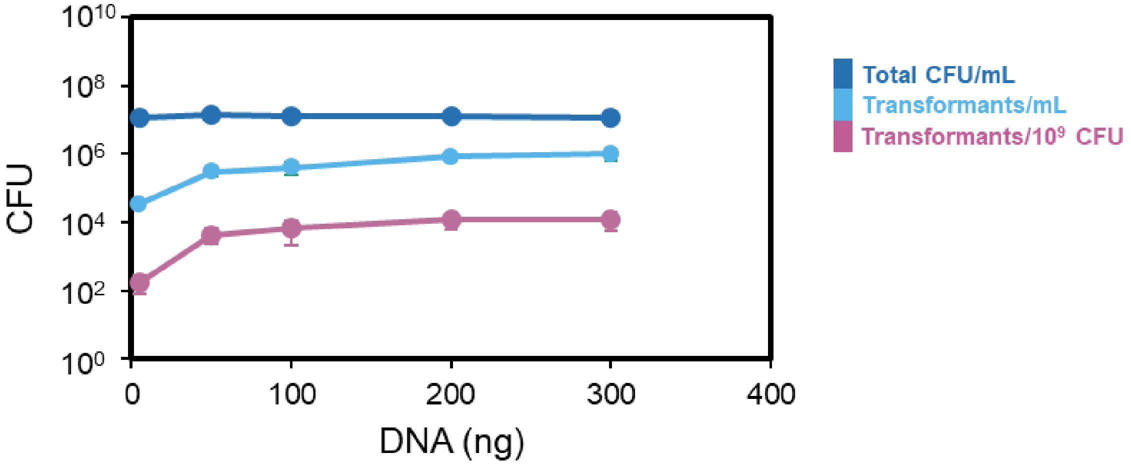


**Figure S6.** Titration of transforming DNA in $P_{xyl}$-*comK* at 1% xylose. The original protocol was used.

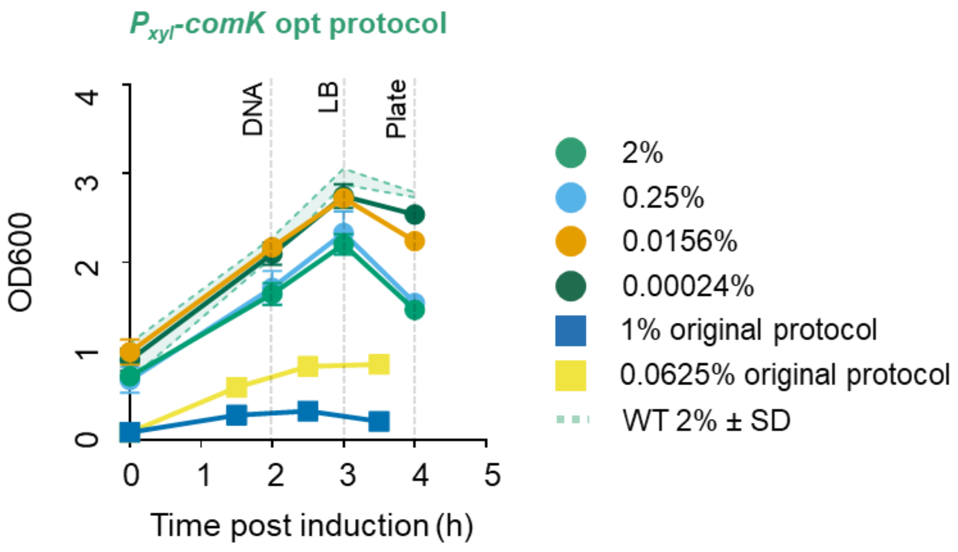


**Figure S7.** $OD_{600}$ growth curves obtained from $P_{xyl}$-*comK* strain under optimized protocol conditions compared to the original protocol at 0.0625% xylose.

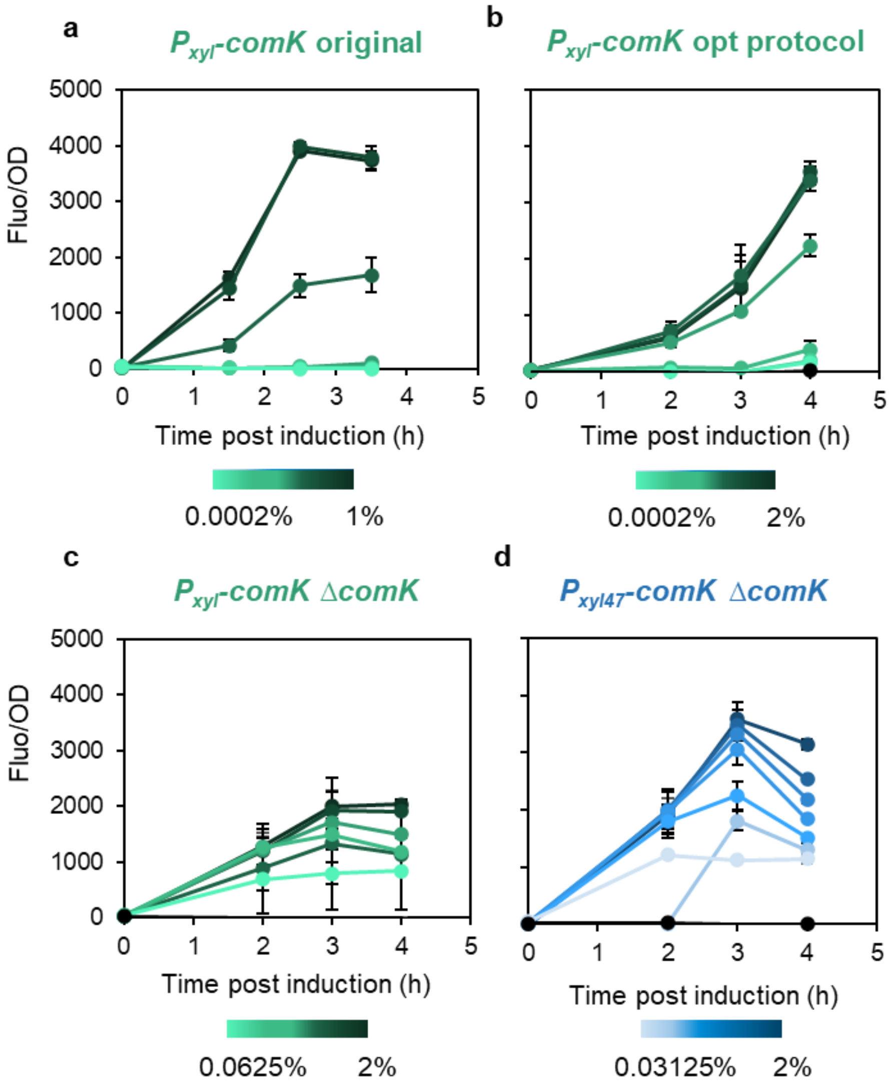


**Figure S8.** Comparison of inducible promoters for competence induction. a) Normalized fluorescence per OD over time for $P_{xyl}$-*comK* under the original protocol, b) $P_{xyl}$-*comK* under the optimized protocol c) $P_{xyl}$-*comK* Δ*comK*, and d) $P_{xyl47}$-*comK* Δ*comK*.

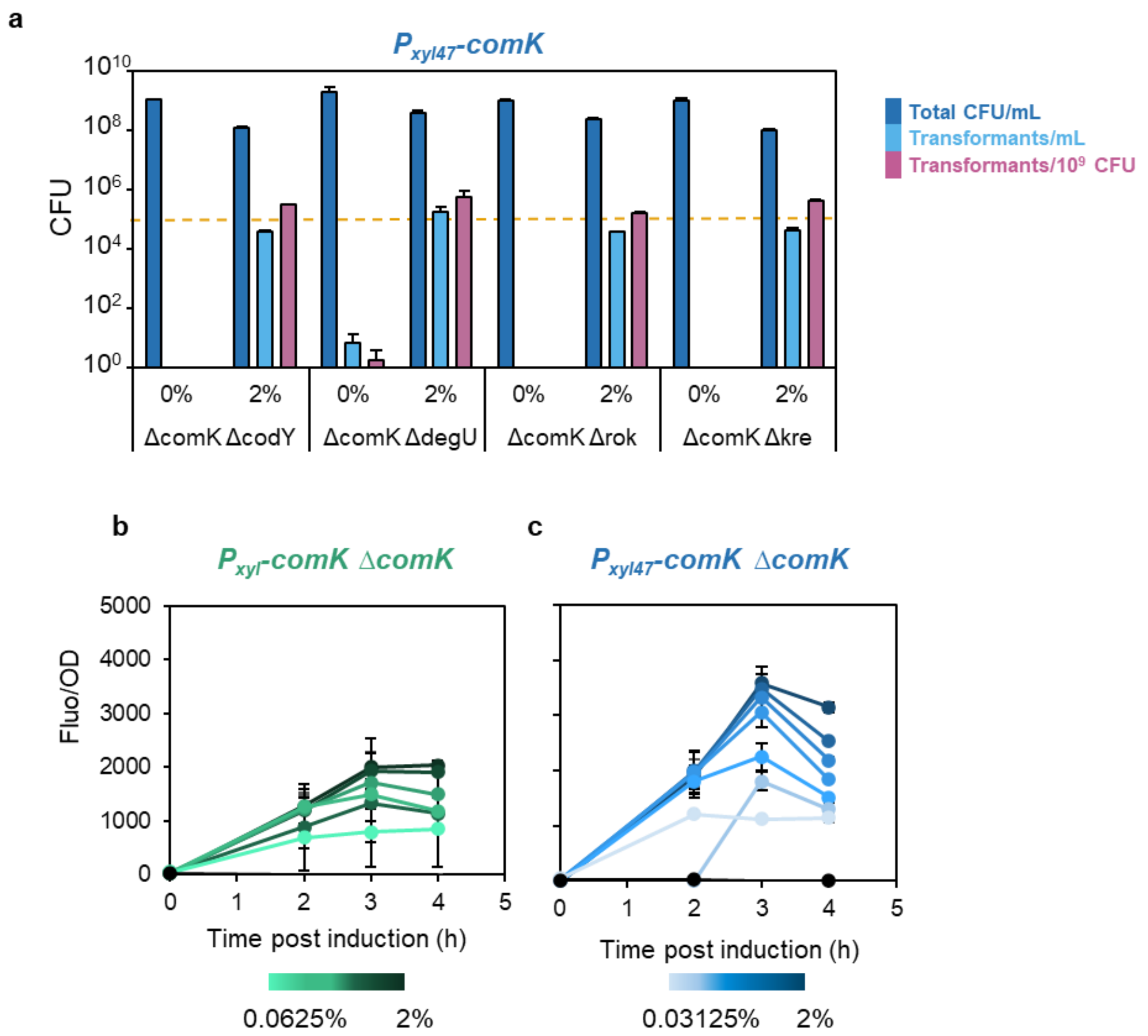


**Figure S9.** a) Effect of knockouts on transformation of $P_{xyl47}$-*comK* Δ*comK* strain. Dashed line represents transformants/mL of $P_{xyl47}$-*comK* Δ*comK* at 2% xylose. b) Normalized fluorescence per OD over time for $P_{xyl}$-*comK* Δ*comK*, and c) $P_{xyl47}$-*comK* Δ*comK*.

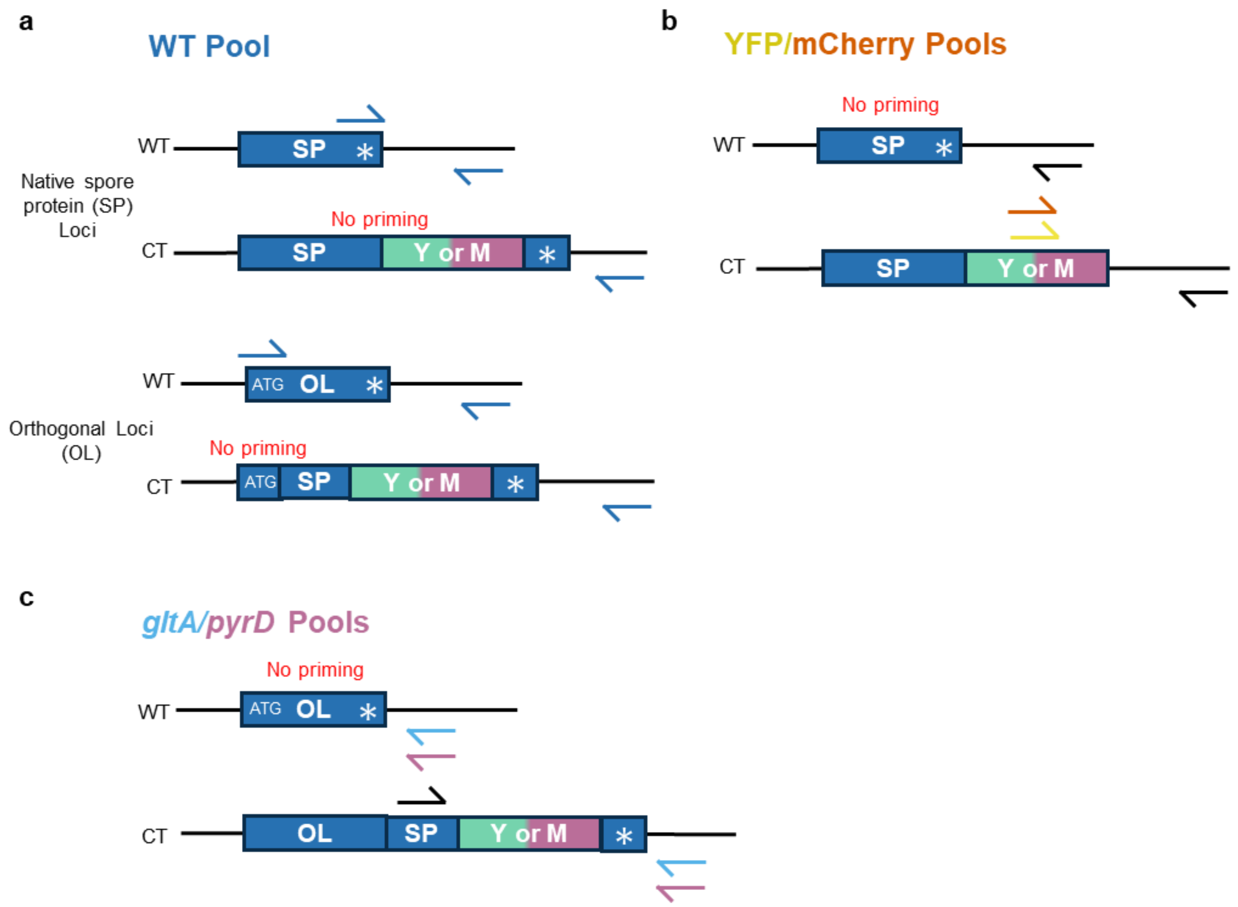


**Figure S10.** MASC PCR assay design. a) Representation of how the WT pool primers bind to WT template vs co-transformed (CT) template at native spore protein (SP) loci and orthogonal loci (OL). Amplification only occurs in the presence of WT template. Primer pairs for each of the seven loci are present in the pool. b) Representation of how the YFP and mCherry pools primers bind to WT vs CT template. The forward primers are specific to either YFP or mCherry and the reverse primers are specific to the locus. Amplification only occurs in the presence of correctly integrated YFP or mCherry. c) Representation of how the *gltA* and *pyrD* pools primers bind to WT and CT template. The forward primers are a pool of five primers for each spore protein. The reverse primer is specific for either *gltA* or *pyrD.* Amplification only occurs in the presence of a spore protein integrated at *gltA* or *pyrD.*

**Table S1.** Plasmids used in this study.

| Plasmid | Description | Source |
| --- | --- | --- |
| pAX01-comK | | *Bacillus* Genetic Stock Center ECE222 |
| pBS1C | | *Bacillus* Genetic Stock Center ECE257 |
| pBS1K | | *Bacillus* Genetic Stock Center ECE731 |
| pDG1730 | | *Bacillus* Genetic Stock Center ECE115 |
| pAX01K-$P_{xyl}$-meYFP | | This study |
| pAX01K-$P_{xyl47}$-meYFP | | This study |
| pAX01K-$P_{mtl}$-meYFP | | This study |
| pAX01K-$P_{hyperspank}$-meYFP | | This study |
| pAX01K-$P_{T7}$-meYFP | | This study |
| pAX01K- $P_{xyl}$-comK | | This study |
| pAX01K-$P_{xyl47}$-comK | | This study |
| pAX01K-$P_{mtl}$-comK | | This study |
| pAX01K-$P_{hyperspank}$-comK | | This study |
| pAX01K-$P_{T7}$-comK | | This study |
| pBS3Klux | | *Bacillus* Genetic Stock Center ECE733 |
| TME2433 | meYFP codon optimized for *B. subtilis* | *Bacillus* Genetic Stock Center ECE752 |
| TME2435 | mCherry codon optimized for *B. subtilis* | *Bacillus* Genetic Stock Center ECE756 |
| pBS3S-$P_{comG}$-meYFP | | This study |
| pBSc241Z | | *Bacillus* Genetic Stock Center ECE711 |
| pENTR-cgeA-C-GusA-His | Native locus integration | Lab stock |
| pENTR-cotG-C-GusA-His | Native locus integration | Lab stock |
| pENTR-cotX-C-GusA-His | Native locus integration | Lab stock |
| pENTR-cotY-C-GusA-His | Native locus integration | Lab stock |
| pENTR-sscA-C-GusA-His | Native locus integration | Lab stock |
| pENTR-cgeA-meYFP | Native locus integration | This study |
| pENTR-cotG-meYFP | Native locus integration | This study |
| pENTR-cotX-meYFP | Native locus integration | This study |
| pENTR-cotY-meYFP | Native locus integration | This study |
| pENTR-sscA-meYFP | Native locus integration | This study |
| pENTR-cgeA-mCherry | Native locus integration | This study |
| pENTR-cotG-mCherry | Native locus integration | This study |
| pENTR-cotX-mCherry | Native locus integration | This study |
| pENTR-cotY-mCherry | Native locus integration | This study |
| pENTR-sscA-mCherry | Native locus integration | This study |

| | | |
|---|---|---|
| pENTR-CT5-cgeA-meYFP | *gltA* integration | This study |
| pENTR-CT5-cotG-meYFP | *gltA* integration | This study |
| pENTR-CT5-cotX-meYFP | *gltA* integration | This study |
| pENTR-CT5-cotY-meYFP | *gltA* integration | This study |
| pENTR-CT5-sscA-meYFP | *gltA* integration | This study |
| pENTR-CT5-cgeA-mCherry | *gltA* integration | This study |
| pENTR-CT5-cotG-mCherry | *gltA* integration | This study |
| pENTR-CT5-cotX-mCherry | *gltA* integration | This study |
| pENTR-CT5-cotY-mCherry | *gltA* integration | This study |
| pENTR-CT5-sscA-mCherry | *gltA* integration | This study |
| pENTR-CT6-cgeA-meYFP | *pyrD* integration | This study |
| pENTR-CT6-cotG-meYFP | *pyrD* integration | This study |
| pENTR-CT6-cotX-meYFP | *pyrD* integration | This study |
| pENTR-CT6-cotY-meYFP | *pyrD* integration | This study |
| pENTR-CT6-sscA-meYFP | *pyrD* integration | This study |
| pENTR-CT6-cgeA-mCherry | *pyrD* integration | This study |
| pENTR-CT6-cotG-mCherry | *pyrD* integration | This study |
| pENTR-CT6-cotX-mCherry | *pyrD* integration | This study |
| pENTR-CT6-cotY-mCherry | *pyrD* integration | This study |
| pENTR-CT6-sscA-mCherry | *pyrD* integration | This study |

**Table S2.** Strains used in this study.

| Strain | Description | Source |
|---|---|---|
| *Bacillus subtilis* 168 | *trpC2* | *Bacillus* Genetic Stock Center 1A1 |
| JL001 | *trpC2;*<br>*lacA::*$P_{xyl}$-*meYFP kmr* | This study |
| JL002 | *trpC2;*<br>*lacA*::$P_{xyl47}$-*meYFP kmr* | This study |
| JL003 | *trpC2;*<br>*lacA::*$P_{mtl}$-*meYFP kmr* | This study |
| JL004 | *trpC2;*<br>*lacA::*$P_{hyperspank}$-*meYFP kmr* | This study |
| JL005 | *trpC2;*<br>*lacA::*$P_{T7}$-*meYFP kmr* | This study |
| JL006 | *trpC2;*<br>*sacA::*P*comG-meYFP spc* | This study |
| JL007 | *trpC2;*<br>*lacA::*$P_{xyl}$-*comK kmr;*<br>*sacA::*P*comG-meYFP spc* | This study |
| JL008 | *trpC2;*<br>*lacA::*$P_{xyl47}$-*comK kmr; sacA::*P*comG-meYFP spc* | This study |
| JL009 | *trpC2;*<br>*lacA::*$P_{mtl}$-*comK kmr;*<br>*sacA::*P*comG-meYFP spc* | This study |
| JL010 | *trpC2;*<br>*lacA::*$P_{hyperspank}$-*comK kmr;*<br>*sacA::*P*comG-meYFP spc* | This study |
| JL011 | *trpC2;*<br>*lacA::*$P_{T7}$-*comK kmr;*<br>*sacA::*P*comG-meYFP spc* | This study |
| BKE16170 | *trpC2;*<br>*codY::erm* | *Bacillus* Genetic Stock Center BKE16170 |
| BKE35490 | *trpC2;*<br>*degU::erm* | *Bacillus* Genetic Stock Center BKE35490 |
| BKE14240 | *trpC2;*<br>*rok::erm* | *Bacillus* Genetic Stock Center BKE14240 |
| BKE06100 | *trpC2;*<br>*kre::erm* | *Bacillus* Genetic Stock Center BKE06100 |
| JL012 | *trpC2;*<br>*lacA::*$P_{xyl}$-*comK kmr;* | This study |

| | | |
|---|---|---|
| | *sacA::*P*comG-meYFP spc*<br>*codY::erm* | |
| JL013 | *trpC2;*<br>*lacA::*$P_{xyl}$-*comK kmr;*<br>*sacA::*P*comG-meYFP spc*<br>*degU::erm* | This study |
| JL014 | *trpC2;*<br>*lacA::*$P_{xyl}$-*comK kmr;*<br>*sacA::*P*comG-meYFP spc*<br>*rok::erm* | This study |
| JL015 | *trpC2;*<br>*lacA::*$P_{xyl}$-*comK kmr;*<br>*sacA::*P*comG-meYFP spc*<br>*kre::erm* | This study |
| BKE10420 | *trpC2;*<br>*comK::erm* | *Bacillus* Genetic Stock Center BKE10420 |
| JL016 | *trpC2;*<br>*lacA::*$P_{xyl}$-*comK kmr;*<br>*sacA::*P*comG-meYFP spc*<br>*comK::erm* | This study |
| JL017 | *trpC2;*<br>*lacA::*$P_{xyl47}$-*comK kmr;*<br>*sacA::*P*comG-meYFP spc*<br>*comK::erm* | This study |
| JL018 | *trpC2;*<br>*lacA::*$P_{xyl}$-*comK kmr;*<br>*sacA::*P*comG-meYFP spc*<br>*comK::erm*<br>*codY::zeo* | This study |
| JL019 | *trpC2;*<br>*lacA::*$P_{xyl}$-*comK kmr;*<br>*sacA::*P*comG-meYFP spc*<br>*comK::erm*<br>*degU::zeo* | This study |
| JL020 | *trpC2;*<br>*lacA::*$P_{xyl}$-*comK kmr;*<br>*sacA::*P*comG-meYFP spc*<br>*comK::erm*<br>*rok::zeo* | This study |
| JL021 | *trpC2;*<br>*lacA::*$P_{xyl}$-*comK kmr;*<br>*sacA::*P*comG-meYFP spc*<br>*comK::erm*<br>*kre::zeo* | This study |
| JL022 | *trpC2;* | |

| | | |
|---|---|---|
| | *lacA::*P$_{xyl}$-*comK kmr;*<br>*amyE*::pBS1C *cmR;*<br>*cgeA::cgeA*-mCherry | |
| JL023 | *trpC2;*<br>*lacA::*P$_{xyl}$-*comK kmr;*<br>*amyE*::pBS1C *cmR;*<br>*cgeA::cgeA*-meYFP | This study |
| JL024 | *trpC2;*<br>*lacA::*P$_{xyl}$-*comK kmr;*<br>*amyE*::pBS1C *cmR;*<br>*cotG::cotG*-mCherry | This study |
| JL025 | *t trpC2;*<br>*lacA::*P$_{xyl}$-*comK kmr;*<br>*amyE*::pBS1C *cmR;*<br>*cotG::cotG*-meYFP | This study |
| JL026 | *trpC2;*<br>*lacA::*P$_{xyl}$-*comK kmr;*<br>*amyE*::pBS1C *cmR;*<br>*cotX::cotX*-mCherry | This study |
| JL027 | *trpC2;*<br>*lacA::*P$_{xyl}$-*comK kmr;*<br>*amyE*::pBS1C *cmR;*<br>*cotX::cotX*-meYFP | This study |
| JL028 | *trpC2;*<br>*lacA::*P$_{xyl}$-*comK kmr;*<br>*amyE*::pBS1C *cmR;*<br>*cotY::cotY*-mCherry | This study |
| JL029 | *trpC2;*<br>*lacA::*P$_{xyl}$-*comK kmr;*<br>*amyE*::pBS1C *cmR;*<br>*cotY::cotY*-meYFP | This study |
| JL030 | *trpC2;*<br>*lacA::*P$_{xyl}$-*comK kmr;*<br>*amyE*::pBS1C *cmR;*<br>*sscA::sscA*-mCherry | This study |
| JL031 | *trpC2;*<br>*lacA::*P$_{xyl}$-*comK kmr;*<br>*amyE*::pBS1C *cmR;*<br>*sscA::sscA*-meYFP | This study |
| JL032 | *trpC2;*<br>*lacA::*P$_{xyl}$-*comK kmr;*<br>*amyE*::pBS1C *cmR;*<br>*gltA::cgeA*-mCherry | This study |
| JL033 | *trpC2;*<br>*lacA::*P$_{xyl}$-*comK kmr;*<br>*amyE*::pBS1C *cmR;* | This study |

| | | |
|---|---|---|
| | *gltA::cgeA*-meYFP | |
| JL034 | *trpC2;*<br>*lacA::*$P_{xyl}$-*comK kmr;*<br>*amyE*::pBS1C *cmR;*<br>*gltA::cotG*-mCherry | This study |
| JL035 | *trpC2;*<br>*lacA::*$P_{xyl}$-*comK kmr;*<br>*amyE*::pBS1C *cmR;*<br>*gltA::cotG*-meYFP | This study |
| JL036 | *trpC2;*<br>*lacA::*$P_{xyl}$-*comK kmr;*<br>*gltA::cotX*-mCherry | This study |
| JL037 | *trpC2;*<br>*lacA::*$P_{xyl}$-*comK kmr;*<br>*amyE*::pBS1C *cmR;*<br>*gltA::cotX*-meYFP | This study |
| JL038 | *trpC2;*<br>*lacA::*$P_{xyl}$-*comK kmr;*<br>*amyE*::pBS1C *cmR;*<br>*gltA::cotY*-mCherry | This study |
| JL039 | *trpC2;*<br>*lacA::*$P_{xyl}$-*comK kmr;*<br>*amyE*::pBS1C *cmR;*<br>*gltA::cotY*-meYFP | This study |
| JL040 | *trpC2;*<br>*lacA::*$P_{xyl}$-*comK kmr;*<br>*amyE*::pBS1C *cmR;*<br>*gltA::sscA*-mCherry | This study |
| JL041 | *trpC2;*<br>*lacA::*$P_{xyl}$-*comK kmr;*<br>*amyE*::pBS1C *cmR;*<br>*gltA::sscA*-meYFP | This study |
| JL042 | *trpC2;*<br>*lacA::*$P_{xyl}$-*comK kmr;*<br>*amyE*::pBS1C *cmR;*<br>*pyrD::cgeA*-mCherry | This study |
| JL043 | *trpC2;*<br>*lacA::*$P_{xyl}$-*comK kmr;*<br>*amyE*::pBS1C *cmR;*<br>*pyrD::cgeA*-meYFP | This study |
| JL044 | *trpC2;*<br>*lacA::*$P_{xyl}$-*comK kmr;*<br>*pyrD::cotG*-mCherry | This study |
| JL045 | *trpC2;*<br>*lacA::*$P_{xyl}$-*comK kmr;*<br>*amyE*::pBS1C *cmR;* | This study |

| | | |
|---|---|---|
| | *pyrD::cotG*-meYFP | |
| JL046 | *trpC2;*<br>*lacA::*P$_{xyl}$-*comK kmr;*<br>*amyE*::pBS1C *cmR;*<br>*pyrD::cotX*-mCherry | This study |
| JL047 | *trpC2;*<br>*lacA::*P$_{xyl}$-*comK kmr;*<br>*amyE*::pBS1C *cmR;*<br>*pyrD::cotX*-meYFP | This study |
| JL048 | *trpC2;*<br>*lacA::*P$_{xyl}$-*comK kmr;*<br>*amyE*::pBS1C *cmR;*<br>*pyrD::cotY*-mCherry | This study |
| JL049 | *trpC2;*<br>*lacA::*P$_{xyl}$-*comK kmr;*<br>*amyE*::pBS1C *cmR;*<br>*pyrD::cotY*-meYFP | This study |
| JL050 | *trpC2;*<br>*lacA::*P$_{xyl}$-*comK kmr;*<br>*amyE*::pBS1C *cmR;*<br>*pyrD::sscA*-mCherry | This study |
| JL051 | *trpC2;*<br>*lacA::*P$_{xyl}$-*comK kmr;*<br>*amyE*::pBS1C *cmR;*<br>*pyrD::sscA*-meYFP | This study |
| JL052 (L24-1) | *trpC2;*<br>*lacA::*P$_{xyl}$-*comK kmr;*<br>*amyE*::pBS1C *cmR;*<br>*cotG::cotG-meYFP;*<br>*gltA::cotG-mCherry* | This study |
| JL053 (L24-2) | *trpC2;*<br>*lacA::*P$_{xyl}$-*comK kmr;*<br>*amyE*::pBS1C *cmR;*<br>*cotG::cotG-meYFP;*<br>*gltA::cgeA-meYFP* | This study |
| JL054 (L24-4) | *trpC2;*<br>*lacA::*P$_{xyl}$-*comK kmr;*<br>*amyE*::pBS1C *cmR;*<br>*cgeA::cgeA-mCherry;*<br>*cotX::cotX-mCherry* | This study |
| JL055 (L24-6) | *trpC2;*<br>*lacA::*P$_{xyl}$-*comK kmr;*<br>*amyE*::pBS1C *cmR;*<br>*gltA::cotX-mCherry;*<br>*pyrD::cotY-mCherry* | This study |
| JL056 (L24-7) | *trpC2;* | This study |

| | | |
|---|---|---|
| | *lacA::*$P_{xyl}$*-comK kmr;*<br>*amyE*::pBS1C *cmR;*<br>*cotY::cotY-mCherry;*<br>*gltA::cgeA-mCherry* | |
| JL057 (L24-8) | *trpC2;*<br>*lacA::*$P_{xyl}$*-comK kmr;*<br>*amyE*::pBS1C *cmR;*<br>*cgeA::cgeA-meYFP;*<br>*cotG::cotG-meYFP;*<br>*gltA::cotG-mCherry* | This study |
| JL058 (L24-9) | *trpC2;*<br>*lacA::*$P_{xyl}$*-comK kmr;*<br>*amyE*::pBS1C *cmR;*<br>*cgeA::cgeA-meYFP;*<br>*cotY::cotY-mCherry;*<br>*gltA::cotG-mCherry* | This study |
| JL059 (L24-11) | *trpC2;*<br>*lacA::*$P_{xyl}$*-comK kmr;*<br>*amyE*::pBS1C *cmR;*<br>*cotX::cotX-meYFP;*<br>*cgeA::cgeA-mCherry;*<br>*gltA::cotY-mCherry* | This study |
| JL060 (L24-12) | *trpC2;*<br>*lacA::*$P_{xyl}$*-comK kmr;*<br>*amyE*::pBS1C *cmR;*<br>*cotY::cotY-mCherry;*<br>*gltA::cotG-meYFP* | This study |
| JL061 (L24-13) | *trpC2;*<br>*lacA::*$P_{xyl}$*-comK kmr;*<br>*amyE*::pBS1C *cmR;*<br>*cotX::cotX-meYFP;*<br>*pyrD::cgeA-mCherry* | This study |
| JL062 (L24-14) | *trpC2;*<br>*lacA::*$P_{xyl}$*-comK kmr;*<br>*amyE*::pBS1C *cmR;*<br>*cotX::cotX-mCherry;*<br>*cotY::cotY-meYFP* | This study |
| JL063 (L24-15) | *trpC2;*<br>*lacA::*$P_{xyl}$*-comK kmr;*<br>*amyE*::pBS1C *cmR;*<br>*cgeA::cgeA-mCherry;*<br>*gltA::cotY-mCherry* | This study |
| JL064 (L24-16) | *trpC2;*<br>*lacA::*$P_{xyl}$*-comK kmr;*<br>*amyE*::pBS1C *cmR;*<br>*cotX::cotX-mCherry;* | This study |

| | | |
|---|---|---|
| | *pyrD::cotY-meYFP* | |
| JL065 (L24-17) | *trpC2;*<br>*lacA::*$P_{xyl}$-*comK kmr;*<br>*amyE*::pBS1C *cmR;*<br>*cgeA::cgeA-meYFP;*<br>*gltA::cotG-meYFP* | This study |
| JL066 (L24-19) | *trpC2;*<br>*lacA::*$P_{xyl}$-*comK kmr;*<br>*amyE*::pBS1C *cmR;*<br>*cotX::cotX-mCherry;*<br>*gltA::cotG-mCherry* | This study |
| JL067 (L24-20) | *trpC2;*<br>*lacA::*$P_{xyl}$-*comK kmr;*<br>*amyE*::pBS1C *cmR;*<br>*cotX::cotX-meYFP;*<br>*gltA::cotY-mCherry* | This study |
| JL068 (L24-21) | *trpC2;*<br>*lacA::*$P_{xyl}$-*comK kmr;*<br>*amyE*::pBS1C *cmR;*<br>*cotY::cotY-mCherry;*<br>*gltA::cotG-meYFP;*<br>*pyrD::cotY-mCherry* | This study |
| JL069 (L24-22) | *trpC2;*<br>*lacA::*$P_{xyl}$-*comK kmr;*<br>*amyE*::pBS1C *cmR;*<br>*cgeA::cgeA-mCherry;*<br>*cotG::cotG-meYFP;*<br>*cotY::cotY-meYFP;*<br>*gltA::cotY-mCherry* | This study |
| JL070 (L24-23) | *trpC2;*<br>*lacA::*$P_{xyl}$-*comK kmr;*<br>*amyE*::pBS1C *cmR;*<br>*cgeA::cgeA-meYFP;*<br>*cotG::cotG-meYFP;*<br>*gltA::cotX-mCherry* | This study |
| JL071 (L24-24) | *trpC2;*<br>*lacA::*$P_{xyl}$-*comK kmr;*<br>*amyE*::pBS1C *cmR;*<br>*cgeA::cgeA-meYFP;*<br>*cotG::cotG-meYFP* | This study |
| JL072 (L24-27) | *trpC2;*<br>*lacA::*$P_{xyl}$-*comK kmr;*<br>*amyE*::pBS1C *cmR;*<br>*cotG::cotG-meYFP;*<br>*cotX::cotX-mCherry;*<br>*gltA::cotY-meYFP* | This study |

| | | |
|---|---|---|
| JL073 (L24-28) | *trpC2;*<br>*lacA::*$P_{xyl}$-*comK kmr;*<br>*amyE*::pBS1C *cmR;*<br>*cotG::cotG-meYFP;*<br>*gltA::cgeA-mCherry* | This study |
| JL074 (L30-2) | *trpC2;*<br>*lacA::*$P_{xyl}$-*comK kmr;*<br>*amyE*::pBS1C *cmR;*<br>*cotX::cotX-mCherry;*<br>*gltA::sscA-meYFP* | This study |
| JL075 (L30-3) | *trpC2;*<br>*lacA::*$P_{xyl}$-*comK kmr;*<br>*amyE*::pBS1C *cmR;*<br>*cotG::cotG-meYFP;*<br>*cotX::cotX-mCherry* | This study |
| JL076 (L30-4) | *trpC2;*<br>*lacA::*$P_{xyl}$-*comK kmr;*<br>*amyE*::pBS1C *cmR;*<br>*cotG::cotG-meYFP;*<br>*sscA::sscA-mCherry* | This study |
| JL077 (L30-7) | *trpC2;*<br>*lacA::*$P_{xyl}$-*comK kmr;*<br>*amyE*::pBS1C *cmR;*<br>*cotY::cotY-mCherry;*<br>*gltA::sscA-mCherry;*<br>*pyrD::cgeA-mCherry* | This study |
| JL078 (L30-8) | *trpC2;*<br>*lacA::*$P_{xyl}$-*comK kmr;*<br>*amyE*::pBS1C *cmR;*<br>*cotG::cotG-meYFP;*<br>*cotX::cotX-mCherry;*<br>*sscA::sscA-mCherry* | This study |
| JL079 (L30-10) | *trpC2;*<br>*lacA::*$P_{xyl}$-*comK kmr;*<br>*amyE*::pBS1C *cmR;*<br>*cgeA::cgeA-meYFP;*<br>*cotY::cotY-mCherry;*<br>*sscA::sscA-meYFP* | This study |
| JL080 (L30-11) | *trpC2;*<br>*lacA::*$P_{xyl}$-*comK kmr;*<br>*amyE*::pBS1C *cmR;*<br>*cotY::cotY-mCherry;*<br>*sscA::sscA-mCherry* | This study |
| JL081 (L30-12) | *trpC2;*<br>*lacA::*$P_{xyl}$-*comK kmr;*<br>*amyE*::pBS1C *cmR;* | This study |

| | | |
|---|---|---|
| | *cotG::cotG-mCherry;*<br>*sscA::sscA-mCherry* | |
| JL082 (L30-13) | *trpC2;*<br>*lacA::*$P_{xyl}$-*comK kmr;*<br>*amyE*::pBS1C *cmR;*<br>*cgeA::cgeA-mCherry;*<br>*cotG::cotG-meYFP;*<br>*cotX::cotX-mCherry* | This study |
| JL083 (L30-14) | *trpC2;*<br>*lacA::*$P_{xyl}$-*comK kmr;*<br>*amyE*::pBS1C *cmR;*<br>*cotG::cotG-mCherry;*<br>*gltA::sscA-meYFP* | This study |
| JL084 (L30-15) | *trpC2;*<br>*lacA::*$P_{xyl}$-*comK kmr;*<br>*amyE*::pBS1C *cmR;*<br>*sscA::sscA-meYFP;*<br>*gltA::cotY-mCherry* | This study |
| JL085 (L30-17) | *trpC2;*<br>*lacA::*$P_{xyl}$-*comK kmr;*<br>*amyE*::pBS1C *cmR;*<br>*sscA::sscA-meYFP;*<br>*gltA::cotG-mCherry* | This study |
| JL086 (L30-18) | *trpC2;*<br>*lacA::*$P_{xyl}$-*comK kmr;*<br>*amyE*::pBS1C *cmR;*<br>*cotG::cotG-meYFP;*<br>*cotY::cotY-mCherry;*<br>*sscA::sscA-mCherry* | This study |
| JL087 (L30-19) | *trpC2;*<br>*lacA::*$P_{xyl}$-*comK kmr;*<br>*amyE*::pBS1C *cmR;*<br>*cotG::cotG-meYFP;*<br>*sscA::sscA-mCherry;*<br>*gltA::sscA-meYFP* | This study |
| JL088 (L30-23) | *trpC2;*<br>*lacA::*$P_{xyl}$-*comK kmr;*<br>*amyE*::pBS1C *cmR;*<br>*cotY::cotY-meYFP;*<br>*sscA::sscA-mCherry* | This study |

**Table S3.** Oligonucleotides used in this study

| Name | Sequence (5' to 3') | Description |
|---|---|---|
| ComK_F1 | GCAGCTGAGCAGCATGTCCAGCC | Amplification of *comK* ermR knockout construct |
| ComK_R1 | GAAGCCTGTCCCTGATTGCGGAG | Amplification of *comK* ermR knockout construct |
| Kre_F1 | GCAGGATGGGCTCCTCATCAATAC | Amplification of *kre* ermR knockout construct |
| Kre_R1 | GGTGTTGGGTGTCAGAGCATC | Amplification of *kre* ermR knockout construct |
| Rok_F1 | CATAATGAAACAAATCTAGAACCATGACG | Amplification of *rok* ermR knockout construct |
| Rok_R1 | CTGTATAAAGCTGCTGCCTTTGAAGATTGC | Amplification of *rok* ermR knockout construct |
| DegU_F1 | CGTCAACTCTCCCTCTTTAATATCACTC | Amplification of *degU* ermR knockout construct |
| DegU_R1 | GCTGGCGGAATTTTATTTCGG | Amplification of *degU* ermR knockout construct |
| CodY_F1 | CATATATGTCTCACCATCCAAGAGC | Amplification of *codY* ermR knockout construct |
| CodY_R1 | CAAACACAGCGTCTTTCGGGAG | Amplification of *codY* ermR knockout construct |
| MASC_WT1_F | CAAAGTAGAAAGCGTTACGTTCTTTTCAtaaCATCC | MASC WT pool, Forward for *cgeA* |
| MASC_WT11_F | GCAATTGCTGGGTAGTCAAAAAGAAATACAAAtaaTCTA | MASC WT pool, Forward for *cotG* |
| MASC_WT25_F | CCATCCTAGTTATCACTCTTGTCCTCtagGA | MASC WT pool, Forward for *cotX* |
| MASC_WT26_F | CACAAAGATAAAAAGCATCATCACAATGGAtaaAC | MASC WT pool, Forward for *cotY* |
| MASC_WT40_F | CGTAGGTGCAGCTTACATCTACtaaGC | MASC WT pool, Forward for *sscA* |
| MASC_WT5_F | GATCGGGGGAGAGGAATTatgACG | MASC WT pool, Forward for *gltA* |
| MASC_WT6_F | GGAGGTGGCGCTGTAatgCTAGA | MASC WT pool, Forward for *pyrD* |
| MASC_1_R1 | CCATCATAAGTGTGAAATCCGGCTGAAACGC | MASC WT, mCherry, and YFP pool, Reverse for *cgeA* |
| MASC_11_R1 | GGCGTCCTTCCGTCCTGTTAACAGAAATAC | MASC WT, mCherry, and YFP pool, Reverse for *cotG* |
| MASC_25_R1 | GCTCTGGTGACAGGCACTGAATCG | MASC WT, mCherry, and YFP pool, Reverse for *cotX* |
| MASC_26_R1 | GAACGTACAGGGGCGTGATTGTCGG | MASC WT, mCherry, and YFP pool, Reverse for *cotY* |
| MASC_40_R1 | CTACCTTTTTAGCTTTTGAGCTGCTGC | MASC WT, mCherry, and YFP pool, Reverse for *sscA* |
| MASC_5_R1 | GGAGATAATACGGCCTTCTTCCAGGTCG | MASC WT, mCherry, and YFP pool, Reverse for *gltA* |
| MASC_6_R1 | CAAGCTTCAAGCACGCTGCCTC | MASC WT pool, Reverse for *pyrD* |

| | | |
|---|---|---|
| MASC_mCherry_F | GCAGAAGGACGCCATTCAACAGG | MASC mCherry pool, Forward for mCherry |
| MASC_YFP_F | GAATTTGTTACAGCAGCAGGCATTACACTG | MASC YFP pool, Forward for YFP |
| MASC_cgeA_F | GCAGTTGGAAATAAAGGGCTTAAAACGGC | MASC SP1 and SP2 pools, Forward for *cgeA* |
| MASC_cotG_F | GCTGTCATAATCATCGTGTCTTTTGTAGTCG | MASC SP1 and SP2 pools, Forward for *cotG* |
| MASC_cotX_F | CTGTTCAATGAGCTGCTCTCTGC | MASC SP1 and SP2 pools, Forward for *cotX* |
| MASC_cotY_F | GGTGGCAAACACTTAAGGTATCGCC | MASC SP1 and SP2 pools, Forward for *cotY* |
| MASC_sscA_F | GAAAGCCACAATATGTTATGATACGCATAGG | MASC SP1 and SP2 pools, Forward for *sscA* |
| MASC_5_R2 | CCTCTATGGTCTAGCTGGCAAAGCATC | MASC SP1 pool, Reverse for *gltA* |
| MASC_6_R2 | CCAGCACCAGTCTCAGCTAC | MASC SP2 pool, Reverse for *pyrD* |
| pENTR_lin_F | ccatcctatggaactgcctcggtgagttttctccttcattacag | Primer for pENTR backbone for amplification of SscA containing constructs |
| pENTR_lin_R | ccgaggcagttccataggatggc | Primer for pENTR backbone for amplification of SscA containing constructs |
| amyE_F | GATGCCAGTGTGTTAGGAACAGATTGG | Primers for amplification of amyE integrated antibiotic resistance constructs from *B. subtilis* genome |
| amyE_R | GCTCAGTGATACCTGCGATCCCTC | Primers for amplification of amyE integrated antibiotic resistance constructs from *B. subtilis* genome |

**DNA construction**

Construction of the five inducible promoter plasmids was completed by PCR (Invitrogen Platinum SuperFi II), gel extraction, and Gibson assembly (NEBuilder HiFi DNA Assembly). The pAX01-comK (BGSC ECE222) plasmid was modified by swapping the erythromycin resistance cassette with a kanamycin resistance cassette from pBS1K (BGSC ECE731). The modified $P_{xyl47}$ promoter was built by removing a vestigial section of the $P_{xyl}$ promoter. The $P_{mtl}$ promoter was introduced to the pAX01K plasmid by amplifying the mannitol promoter from *B. subtilis*. The $P_{hyperspank}$ promoter was amplified from a gene fragment (Twist Biosciences). A gene fragment containing lacI was also built into the pAX01K-$P_{hyperspank}$-comK and pAX01K-$P_{hyperspank}$-YFP plasmids. The PT7 containing plasmids were assembled in 5 fragments: the vector backbone, a gene fragment containing $P_{hyperspank}$-T7RNAP a terminator from pDG1730 comK from pAX01K and lacI from a gene fragment. To build the reporter strains, *B. subtilis* codon optimized meYFP was obtained from BGSC ECE752.

To build the $P_{comG}$-YFP competence reporter, the reporter was first cloned in pDG1730. meYFP was amplified from BGSC ECE752 and $P_{comG}$ was amplified from the *B. subtilis* genome. The entire assembled cassette was then amplified and assembled with pBS3K.
The erythromycin resistance containing knockout cassettes for *comK, kre, rok, degU,* and *codY* were amplified with 2 kb homology arms from BGSC strains BKE10420, BKE06100, BKE14240, BKE35490, BKE16170 using oligos indicated in Supplementary Table 3. The zeocin resistance containing knockout cassettes for *kre, rok, degU,* and *codY* were assembled by overlap extension PCR using the homology arms from the erythromycin knockout cassettes and the zeocin resistance maker from pBSc241Z (BGSC ECE711).

The pENTR co-transformation constructs were cloned from lab stocks of pENTR plasmids for integration of GUS fusions at *cgeA, cotG, cotX, cotY*, and *sscA*. The GUS CDS was removed and replaced with *B. subtilis codon* optimized mCherry and meYFP from TME2433 and TME2435. To build the co-transformation constructs for integration at *gltA* and *pyrD*, 2 kb upstream and downstream homology arms for the orthogonal sites replaced those for native sites. Since SscA containing constructs were not stable in *E. coli*, once finished plasmids were cloned, the DNA was amplified with pENTR_lin_F and pENTR_lin_R for co-transformation. The *amyE* chloramphenicol, spectinomycin, and erythromycin antibiotic integration cassettes are derived from the antibiotic resistance cassettes from pBS1C, pBS3Klux, and pBS1E. Two kb homology arms from *B. subtilis* amyE were added by overlap extension PCR and amplified with amyE_F and amyE_R.
